# Giotto Suite: a multi-scale and technology-agnostic spatial multi-omics analysis ecosystem

**DOI:** 10.1101/2023.11.26.568752

**Authors:** Jiaji George Chen, Joselyn Cristina Chávez-Fuentes, Matthew O’Brien, Junxiang Xu, Edward Ruiz, Wen Wang, Iqra Amin, Irzam Sarfraz, Pratishtha Guckhool, Adriana Sistig, Guo-Cheng Yuan, Ruben Dries

## Abstract

Emerging spatial omics technologies continue to advance the molecular mapping of tissue architecture and the investigation of gene regulation and cellular crosstalk, which in turn provide new mechanistic insights into a wide range of biological processes and diseases. Such technologies provide an increasingly large amount of information content at multiple spatial scales. However, representing and harmonizing diverse spatial datasets efficiently, including combining multiple modalities or spatial scales in a scalable and flexible manner, remains a substantial challenge. Here, we present Giotto Suite, a suite of open-source software packages that underlies a fully modular and integrated spatial data analysis toolbox. At its core, Giotto Suite is centered around an innovative and technology-agnostic data framework embedded in the R software environment, which allows the representation and integration of virtually any type of spatial omics data at any spatial resolution. In addition, Giotto Suite provides both scalable and extensible end-to-end solutions for data analysis, integration, and visualization. Giotto Suite integrates molecular, morphology, spatial, and annotated feature information to create a responsive and flexible workflow for multi-scale, multi-omic data analyses, as demonstrated here by applications to several state-of-the-art spatial technologies. Furthermore, Giotto Suite builds upon interoperable interfaces and data structures that bridge the established fields of genomics and spatial data science, thereby enabling independent developers to create custom-engineered pipelines. As such, Giotto Suite creates an immersive ecosystem for spatial multi-omic data analysis.

## Introduction

Biological tissues are organized in a hierarchical and structured manner that is optimized for their specific functions and occur at various scales. Examples are abundant, including the subcellular localization of transcripts across the polarized axis of intestinal enterocytes^1^, the different roles for individual liver cells dictated by hepatocyte zonation^2^, multi-cellular niches comprised of macrophages, endothelial and cancer cells that promote metastasis^3^, or the layered organization of the brain^4^. As a result, large-scale efforts to systematically create spatial maps from various tissues^5^ and disease states^6^ are occurring with increased frequency. With the introduction and advancement of spatial omics technologies, including spatial transcriptomics^7–10^, proteomics^11–14^, and others^15^, researchers have the ability to visualize and analyze various molecular analytes within their tissue context and across different levels of the biological organization. Each of these techniques provides a unique level of resolution and data modality, creating an intricate, multidimensional map of biological tissue. Regardless of the scale, the individual units of a tissue are single cells whose phenotypes are defined by the interplay of multiple regulatory layers, such as variations in the (epi-)genome, transcriptome, proteome, and metabolome. Accumulating research highlights the importance of integrating both multi-omics data within single cells and multi-scale patterns within the tissue. Connecting both the intrinsic and extrinsic layers of variation will provide the foundation to understand how the activities of single cells jointly coordinate tissue function and organization in a systems biology manner.

For instance, while spatial transcriptomics allows the exploration of gene expression in a spatial context, spatial proteomics on the same - or immediately adjacent tissue slice - provides complementary insights into the spatial distribution of proteins. Furthermore, most technologies can be combined with imaging, which captures tissue and cellular morphology. Similarly, multiple serial sections from a tissue of interest can be profiled to create a 3-dimensional (3D) representation^16–18^. Hence, by integrating these multi-scale and multi-modal data, a comprehensive systems biology perspective of biological tissue can be attained, offering an unparalleled, two-or three-dimensional view of biology *in vivo*. These integrative approaches promise to significantly advance our understanding of the complex spatial and functional relationships among cells and tissues, leading to a more precise and nuanced understanding of biological processes in health and disease. However, most software engineering approaches are not purposefully designed to fully capture the increasing complexity associated with multi-scale and multi-modal datasets in a technology-agnostic manner. Furthermore, tools and methods to access and work with the full breadth of information that is available within these emerging spatial datasets is either performed in isolated environments or completely lacking. Hence, there is a dire need for increased method and data engineering development along with the spatial technologies and datasets.

The programming language R is widely used for analyzing biological data. It has seen a strong increase in users across the fields of biomedical sciences since it offers an array of tools for statistical and genomics analysis. For instance, the Bioconductor project has a central role in more advanced omics analysis and has created a collaborative community for method development and interoperability^19^. In parallel, the geospatial field has a long history of working with various spatial data, including raster and vector-based data types^20^. However, implementations of the many associated and established spatial data analysis or simulation methods are rare within the field of biomedical sciences. Here we leveraged our previous expertise from Giotto^21,22^ to create a rich and inclusive ecosystem for spatial data analysis and engineering. This ecosystem includes an innovative and immersive spatial multi-modal data framework, called Giotto Suite, and modules to represent, analyze, and visualize any type of spatial omics data. In addition, it maximizes and connects with other existing exploratory data ecosystems in genomics and spatial data science to provide users with easy workflows for a plethora of spatial downstream analyses. Finally, it facilitates the building of novel methods and applications by external developers that are accessible to large user groups.

## Results

### Giotto Suite core framework

Giotto Suite is a new modular suite of R packages that together create a holistic spatial data analysis ecosystem (**Fig. 1A**). It is technology-agnostic and designed to work with the ever-increasing size and complexity of spatial datasets, including innovative implementations for multi-modal and multi-resolution dataset representation and integration. It couples easy-to-use workflows on a large variety of spatial technologies with extended documentation, including examples for numerous downstream spatial analysis methods and visualizations (**Supplementary Fig. 1A**). Importantly, Giotto Suite has been specifically designed to underscore the FAIR principles and promote community building in an open-source software environment^23^.

**Figure 1:**
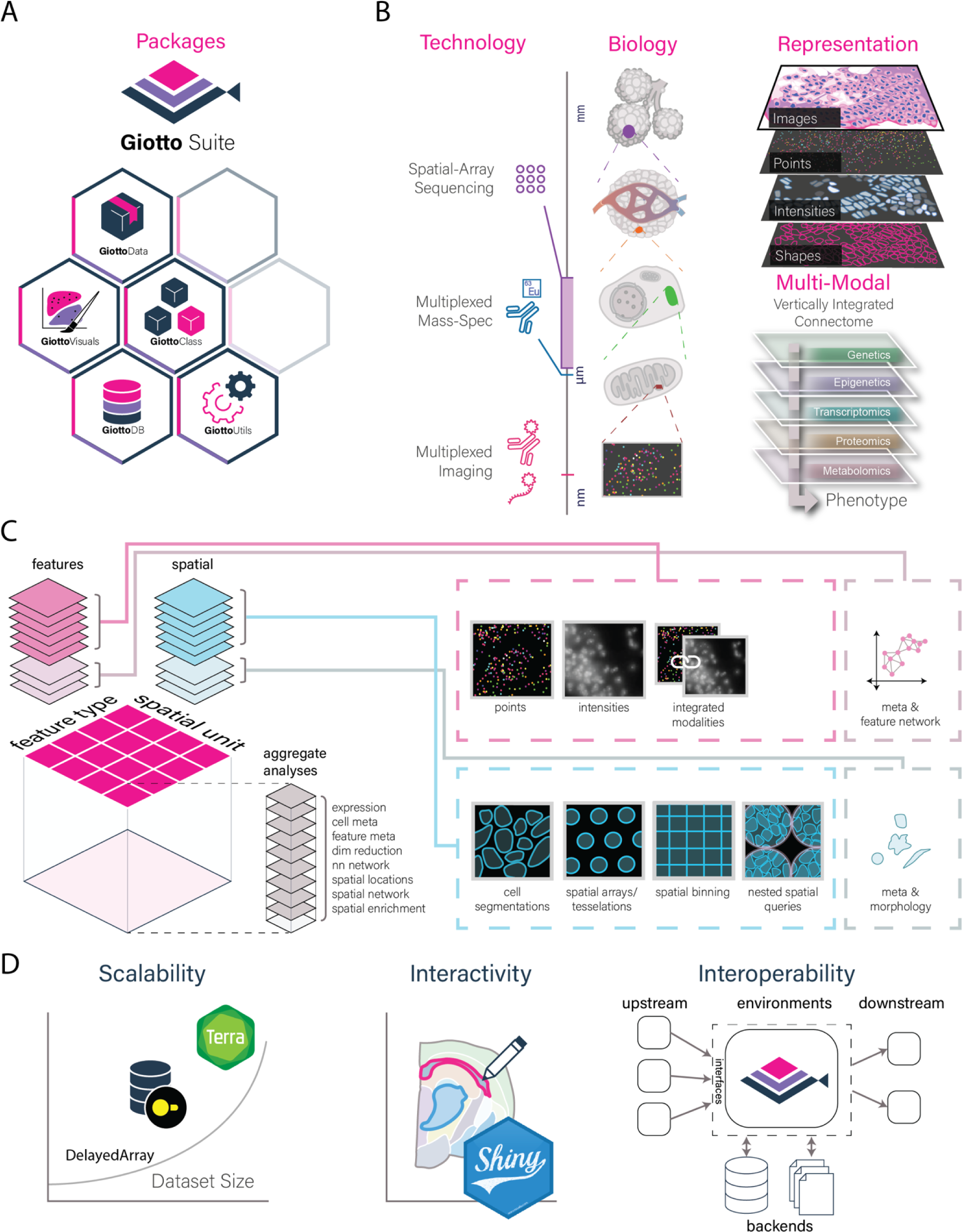
Giotto Suite ecosystem and core data framework. **A)** Giotto Suite consists of a suite of integrated R packages supported by a technology-agonistic core data framework. **B)** Illustration depicting the conceptual and flexible integration of spatial datasets across multiple length scales and data modalities in Giotto Suite. **C)** Schematic depicting how Giotto Suite’s core framework represents diverse spatial datasets by features (molecular analytes) and spatial units (e.g. cell or grid) thereby facilitating various multi-scale and multi-omics data integration. **D)** Pictograms summarizing key implementations for flexibility, scalability, interactivity, and interoperability.

At the core of Giotto Suite is an innovative data framework that is designed to be agnostic to spatial omics technology and to maximally capture and represent the multi-scale and multi-modal biological data (**Fig. 1B**). This framework is designed based on two core principles. First, any type of spatial data can be efficiently used and represented by dedicated data structures, which are built on top of classes within the geospatial *terra* package and emphasize retaining raw information to a maximal degree. These structures include `giottoPoints, giottoPolygon, and giottoLargeImage` that capture respectively point, vector-based, and image or raster-like information (**Fig. 1B** and **Supplementary Fig. 1B**). `giottoPoints` represents points information (e.g. individual transcripts) and their associated spatial coordinates generated by multiplexed *in situ* hybridization techniques, thus ensuring that subcellular resolution is retained. Similarly, it can also be used to work with sequencing-based methods that generate spatial data from individual arrays with dimensions at the subcellular scale^9,24^. `giottoPolygon` is a versatile class used to represent the spatial organization and structures of biological shapes or annotated regions, including biological segmentation results (e.g. cell or organelle boundaries), flexible spatial arrays, or uniform grid and tessellation structures. Hence, it can be used to represent several types of information, ranging from (sub)cellular or tissue structures to external biological information (e.g. pathology annotations). Finally, almost all spatial technologies include either an associated tissue image (e.g. H&E image) or build a dataset through sequential imaging (e.g. protein intensities for cyclic antibody multiplexing technologies^12,25^), and this type of data is represented with `giottoLargeImages`. As such, it provides a dual role by allowing both efficient visualization and extraction of the data.

The second principle focuses on data organization. Giotto Suite implements an approach that can be easily modified to allow the integration of multiple datasets with different modalities and/or resolutions (**Fig. 1C**). A key step herein is to organize data types based on their feature type and spatial unit (**Supplementary Fig. 1C**). The feature type refers to the corresponding data modalities, whereas the spatial unit refers to the underlying spatial structure (e.g. nucleus, cell, or abstract grid/spot) and is represented by the `giottoPolygon` class. In this manner, an unlimited number of feature type-spatial unit aggregations can be performed and efficiently stored for further downstream specialized integration methods (**Supplementary Fig. 2A, B**). In addition, provenance and hierarchy can be encoded within the aggregation or integration steps to build a custom hierarchical tissue structural model. Hence, the core framework in Giotto Suite facilitates the aggregation or integration of multiple different feature types at multiple spatial units at both technical and biological levels.

To support the core framework the original and convoluted S4 Giotto class has been redesigned and supplanted by a light-weight design that underscores the generation of independent S4 subclasses representing different data types and structures. These subclasses are both extensible and easy to maintain. This new design underscores a commitment to good practice object-oriented programming (OOP) principles, providing a platform with increased flexibility for future tooling, visualization, and framework development. Additionally, this design is oriented toward ensuring backward compatibility, thereby creating a seamless experience for long-term users and adopters. Moreover, improved and unified accessor and show functions were created as well as *terra* spatial generics for each subobject. Continuous integration and unit testing have been added to facilitate contributions from external developers. Finally, significant efforts to improve scalability, interactivity, and interoperability (**Fig. 1D**) were implemented to augment the new core framework from Giotto Suite as demonstrated through the following vignettes.

### Vignettes for multi-scale and multi-omic analyses

To demonstrate the ability of Giotto Suite and highlight its spatial technology-agnostic (**Supplementary Fig. 3**), flexible, and scalable implementations, we showcase various applications on specific datasets generated by some of the latest spatial technologies as follows.

*Multiscale and expansive framework.* Biological processes occur at multiple scales and resolutions (**Fig. 2A**) and will lead to different - but related - scientific questions throughout the anatomical hierarchy of the tissue. Giotto Suite’s core framework (**Fig. 1C**) facilitates joint data representation and analysis at any level (**Fig. 2A & Supplemental Fig. 4A-B**). To demonstrate this utility, we use a subset of the MERFISH FFPE Human breast cancer dataset. First, tissue structures are annotated with increasing granularity, such as tissue domains, niches, individual cell types, or nuclei (**Fig. 2B & Supplemental Fig. 4B**). Next, the data organization facilitates efficient queries between different spatial scales and can also be used to assess how independent clustering results at different scales (e.g. nuclei vs cells) are related (**Supplemental Fig. 4C-E**).

**Figure 2:**
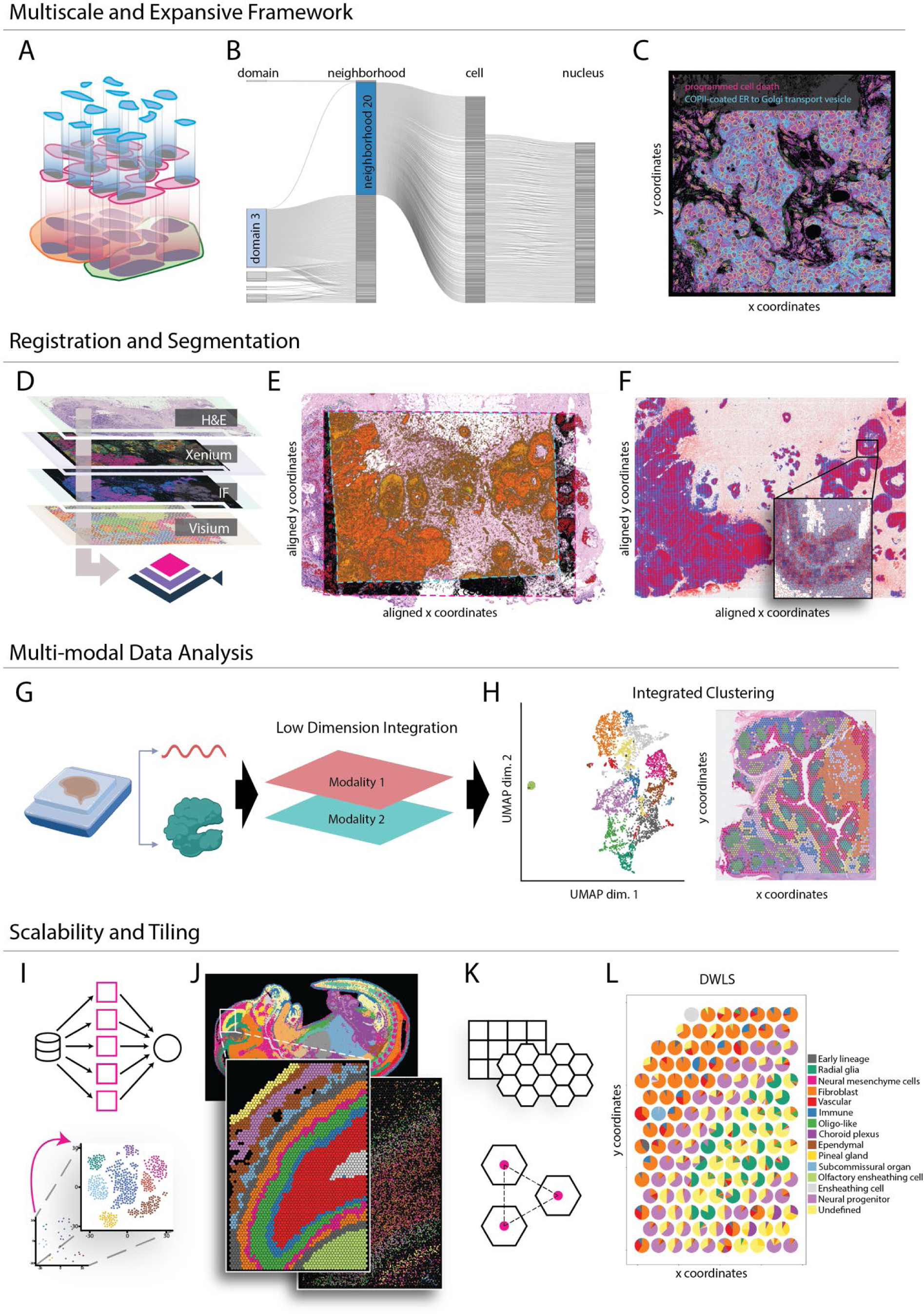
Representative vignettes implemented in Giotto Suite. **A-C)** Multi-scale analysis. **A)** Pictogram depicting the hierarchy of multiple biological scales. **B)** Sankeyplot showing the hierarchical relationships between different spatial units in Giotto Suite. **C)** Overlay of multi-scale annotations of a MERFISH dataset; depicting nuclear (magenta) and cytoplasmic (cyan) enriched transcripts for all genes in GSEA terms related to “programmed cell death” and “COPII-coated ER to Golgi transport vesicle”, respectively. The nuclear boundary is defined with a yellow border, and the cell body is filled in with dark green. **D-F)** Registration and segmentation. **D)** Pictogram showing multi-modal co-registration from serial tissue sections. **E)** Aligned Xenium (Black), Visium (Orange), and immunofluorescence (Red and Green) spatial datasets from a single breast cancer sample (10X Genomics). **F)** Overlay of HER2 protein (rasterized intensity, blue gradient) and ERBB2 transcripts (points, red) after co-registration, inset shows zoomed-in region. **G-H**) Multi-modal data analysis. **G)** Pictogram depicting multi-modal data analysis from two different modalities obtained from the same tissue slice. **H)** Integrated clustering results (RNA + Protein) from the 10X Genomics CytAassist Visium Human Tonsil dataset. **I-L)** Scalable data analysis. **I)** Pictograms depicting on-disk backends, parallelization, and projection to improve scalability within Giotto Suite. **J)** Stereo-seq datasets processed at bin1 and analyzed through multiple levels of hexagon aggregation, the inset depicts a zoomed-in subset and visualization of niche clustering on Leiden annotations and associated selected transcripts with random jitter. **K-L)** Tiling and pseudo-spatial data creation **K)** Schematic pictograms for different tiling options and tunable parameters. **L)** Spatial deconvolution plot showing the percentage of individual brain cell types for the pseudo-visium dataset generated from the stereo-seq subset in J.

In addition to common analysis pipelines that typically treat each cell as the basic unit, Giotto Suite’s framework allows users to carry out subcellular analysis as well. Individual transcript locations can be queried against any pre-defined spatial units and used to identify genes or gene sets that are spatially enriched at subcellular organelles, such as nucleus vs cytoplasm (**Fig. 2C** & **Supplementary Fig. 5A,B**) or detect transcripts that are found preferentially inside or outside cell boundaries (**Supplementary. Fig. 5C**). For example, Gene Set Enrichment Analysis (GSEA)^26^ identified enrichment of (ribo-)nucleotide related genes within cells while, interestingly, genes associated with the extracellular matrix are also often located outside cells themselves (**Supplementary Fig. 5D,F**). When comparing transcript location preferences inside the nucleus or cytoplasm, genes associated with apoptosis were enriched within the nucleus, and genes linked to cytoplasmic membrane structures (i.e. Golgi transport and endoplasmic reticulum) were found mostly within the cytoplasm (**Supplementary Fig. 5E,G**). Finally, the Giotto Suite framework facilitates subcellular 3D data analysis. To demonstrate this workflow, we use a subset of the MERFISH mouse brain dataset (v1.0, May 2021) as an example. Within this dataset individual transcripts are detected in seven adjacent z-stacks, each separated by approximately 1.5 µm (**Supplementary Fig. 6A**). In addition, each z-stack also contains its own cell polygon information, such that transcripts can be accurately assigned to the correct cell polygon (**Supplementary Fig. 6A,B**). Users can efficiently aggregate the multiple stacks to create spatial networks at the transcript level (**Supplementary Fig. 6B**) or use layer-specific information to assess technological or biological differences between layers (**Supplementary Fig. 6C-H**).

*Registration and segmentation.* Spatial omics assays can profile different molecular analytes in thin tissue sections. The use of multiple adjacent tissue sections has become a popular strategy to create 3D spatial datasets^16–18,27^ or to integrate multiple technologies^28–30^ that are not directly compatible on the same section slice (**Fig. 2D**). This approach provides individual labs, or larger consortia with multiple groups, the opportunity to create more complex datasets that can unravel the intricacies of cellular decision making in the context of tissue architecture. Tools for aligning or co-registering one or more tissue slices have been developed previously^31,32^ or are currently being adapted for spatial biology purposes^33,34^. However, capturing and working with different spatially aligned datasets, each with its own resolution and data output, is still challenging. The Giotto Suite framework is uniquely suited to work with multiple co-registered spatial datasets due to its flexible handling of different scales and data types. To demonstrate this capacity, we use the recent spatial multi-modal breast cancer dataset from 10X Genomics, consisting of Visium, Xenium, H&E, and immunofluorescence (IF) as a concrete example^29^. First, tissue registration was performed by STalign^34^ and the transformed spatial datasets, including Xenium, Visium, IF and H&E images, were subsequently used to create a multi-modal and multi-scale Giotto object for downstream analysis, including RNA, protein, and morphology information (**Fig. 2E** & **Supplementary Fig. 7A**). Next, correlations between corresponding spatial protein and RNA levels, such as for the B-cell (CD20 and *MS41*) and breast cancer (HER2 and *ERBB2*) markers demonstrated moderate to high correlations between the protein and transcript analytes, respectively r = 0.21 and 0.60 (**Fig. 2F & Supplementary Fig. 7B,C**). Similarly, systematic comparisons were performed for transcript counts from genes present in both the Visium (Sequencing) and Xenium (*in situ* hybridization) datasets (**Supplementary Fig. 7D**). This analysis indicated that on average concordance between both technologies is good (median r = 0.42) and that lower correlations are associated with low detection levels (**Supplementary Fig. 7E**). For example, the high expressed FASN gene showed a similar spatial expression pattern for both Visium and Xenium (**Supplementary Fig. 7F**) however, the opposite was seen for the low expressed HDC gene (**Supplementary Fig. 7G**).

Another key task for many new spatial technologies is cell segmentation and subsequent transcript aggregation. Giotto Suite can read and simultaneously store various segmentation output formats, including mask files or geojson files, and this capacity was illustrated for several common methods^35–38^ (**Supplementary Fig. 8A**). To assess how segmentation choice might lead to variable outcomes in cell type composition two complementary analyses were performed. First, a Giotto object was created containing both original segmented cells, i.e. provided by 10X Genomics, and cells segmented by Baysor^36^, which also uses transcript density. Next, transcripts were aggregated within cells from both segmentation results and co-clustered to obtain joint annotations for both methods (**Supplementary Fig. 8B,C**). The results did not only show a difference in the number (Baysor: 220,845 vs original: 164,471) and pixel area size (Baysor: 4.9 vs original: 14.2) (**Supplementary Fig. 8D**) of detected cells, but also striking differences in cellular composition, such as for stromal cells (**Supplementary Fig. 8B,E,F**) which are key players in breast cancer progression^39,40^. Next, a more controlled analysis was performed by resizing (i.e. 25% downscaled) cell segmentation results from Imaging Mass Cytometry (IMC) data^41^ (**methods** & **Supplementary Fig. 9A**). Unbiased clustering with k-means (k = 7) of both the original and downscaled cells showed a substantial number (26%) of cell annotation class switches that are dispersed across the region of interest (**Supplementary Fig. 9B-E**). Together, these observations highlight how Giotto Suite provides a standardized ecosystem to study how segmentation choices or strategies can affect downstream spatial analysis.

*Multi-modal data analysis.* Emerging technologies provide a great opportunity to investigate the relationship between different data modalities. A major addition to Giotto Suite is the support of multi-modal data analyses and integration to obtain a more comprehensive characterization of cell states. This is made possible by the addition of specific features facilitated by the core framework. First, multiple data modalities (e.g. RNA and Protein) can be stored at the same spatial locations (**Fig. 2G**). Here the multi-omic data may either be obtained from the same cells or assembled from different cells through image alignment, as described previously (**Fig. 2D**). Next, efficient structures for combining and re-weighting graph structures from different modalities have been implemented that could support various statistical integration approaches^42–44^ (**Supplementary Fig. 10A**). To demonstrate how Giotto Suite handles multi-modal data, we analyzed the Visium CytAssist human tonsil dataset (10X Genomics), containing multi-omics information for RNA and surface proteins (**Supplementary Fig. 10B-C**). By analyzing each data modality separately, we identified 12 cell clusters from RNA and protein profiles, respectively (**Supplementary Fig. 10B**). The difference between the clustering results suggests these data modalities contain complementary information. By using a weighted nearest-neighbor (WNN) method^43^, both RNA and protein information were integrated thereby obtaining 13 integrated cell clusters (**Fig. 2H**). Compared to the outcome from individual modalities, the multi-omic clustering result performs better and its spatial pattern clearly resembles the histological structures in the tissue image (**Fig. 2H& Supplementary Fig. 10C**). This is further corroborated by the spatial distribution of different cell types, which were estimated from a cell-type deconvolution algorithm^45^. The resulting pattern follows the known architecture of the tonsil (**Supplementary Fig. 10D**).

*Scalability and tiling.* Advancements in spatial technologies have led to ever-growing datasets that can be stored in large databases^46–48^ and can easily contain spatial and expression information from millions of cells. Even high-performance computing infrastructure can struggle to process these datasets with conventional methods. To alleviate the challenge of scalable analysis, several complementary tools are implemented in Giotto Suite, including optimized parallel coding, delayed on-disk calculations, and data projection strategies (**Fig. 2I, Supplementary information**). The Giotto Suite package is also built, tested, and available on terra.bio as a cloud-based solution to accommodate users who have no access to high-performance computing infrastructure or would like additional scaling (**Supplementary Fig. 11A**).

As an illustrating example, we analyzed a stereo-seq dataset obtained from a mouse embryo at embryonic day 16.5 (“E16.5_E2S6”) from the Mouse Organogenesis Spatiotemporal Transcriptomic Atlas (MOSTA)^24^. We focused on a whole sagittal section at its highest resolution (bin1) (**Fig. 2J**). The dataset contains 378 million transcripts from 292 million bins covering the whole transcriptome. Storing the raw cell gene matrix alone takes about 40GB of memory. To facilitate working with large spatial data, a database backend was implemented to provide on-disk S4 representations for points and polygons information as `dbPointsProxy` and `dbPolygonProxy`, respectively (**Supplementary Fig. 11B**). These representations respond to spatial manipulation generics and can also be directly converted into corresponding *terra* objects that are native to the Giotto Suite framework. In order to increase responsiveness and allow in-memory operations, lazy evaluation is used when possible, and spatial chunking is implemented on these S4 classes. To support interactive usage of these objects, they are also implemented with high-performing plotting methods. Finally, any resulting large aggregated expression matrices are handled by using a delayed HDF5matrix^49^ on-disk backend (**Supplementary Fig. 11C**). Users can simultaneously achieve significant additional computing speed gains by using data projection strategies for initial data exploration. Similar to a standard exploratory data analysis pipeline, it facilitates the optimization of parameters in a computation-efficient and more responsive manner.

In parallel, Giotto Suite also offers flexible tiling and tessellations (**Fig. 2K**) that can be interpreted by the spatial framework layer (**Fig. 1C** & **Supplementary Fig. 2A**). Tiling is a popular strategy to analyze large-scale data at multiple scales or resolutions (**Fig. 2J & Supplementary Fig. 12A-C**) but can also be used to create pseudo-datasets with a custom grid configuration to support benchmark analyses. For example, a more granular ‘pseudo-visium’ dataset was created from a brain region in the stereo-seq mouse embryo dataset (**Supplementary Fig. 12D**). Next, spatialDWLS was used to deconvolve each pseudo-visium spot and to assess deconvolution accuracy relative to the original and ground-truth stereo-seq data (**Fig. 2L** & **Supplementary Fig. 12D**). Together, tiling and scalability implementations provide users with the tools needed to analyze large-scale data at custom resolutions or patterns.

*Interactivity & Interoperability.* To aid users in their exploration of the relationship between molecular, histomorphological, and pathological changes within a tissue, we have created an integrated and interactive Shiny app for refined annotation and region selection in Giotto Suite (**Supplementary Fig. 13A**). This tool allows users to manually annotate multiple spatial regions using an html widget. Notably, information within the selected regions (e.g. cell identities) is immediately available in the Giotto object and can thus be directly used in any other downstream analyses (e.g. differential expression) (**Supplementary Fig. 13B**). Additionally, we have developed an interactive tool to plot and subset 3D datasets (e.g. mouse brain with MERFISH^27^) that facilitate subsetting slices in the z-axis and select cell types or clusters. The subsetted cells can be used for downstream analysis such as the comparison of gene expression patterns across diverse slices (**Supplementary Fig. 13C**).

During the past years, many groups have developed innovative tools and methods for spatial transcriptomics data analysis^50,51^. Giotto Suite provides several utilities to facilitate interoperability with these external tools, including rich data structures, accessor functions, and plugins. Giotto Suite also provides built-in converter functionality for the R/Bioconductor *SpatialExperiment*^52^, python AnnData^53^, and Seurat^54^ classes, and classes used in the open spatial sciences field within R^55^. For example, Giotto Suite users can use the bi-directional converters (**Supplementary Fig. 14A, B, C**) to effortlessly use tools developed with Seurat or in Bioconductor and subsequently combine and visualize the final results within the Giotto framework. Similarly, the Giotto to AnnData class converter functions can be used to download and access all datasets within the Spatial Omics DataBase (SODB)^46^ (**Supplementary Fig. 14B**). In addition, a modified AnnData version for the Bento pipeline is also available and allows users to perform various RNA localization pattern analyses (**Supplementary Fig. 14D-F**). Finally, various classes (e.g. *terra, sf, stars,* etc.) and associated methods and statistics used in the R spatial sciences field are easily accessible (**Supplementary Fig. 15A**) and can be directly created from Giotto (sub-)classes through “as’’ converter functions (**Supplementary Fig. 15B**). In this manner, other methods and packages can utilize the accessor functions in Giotto Suite to extract information from individual Giotto Suite slots. For example, interpolation methods such as kriging in the *gstat* package, can be easily combined with the Giotto Suite framework to develop unique ways to enhance low-resolution spatial datasets (**Supplementary Fig. 15C**). Altogether, close integration with these other large and established ecosystems significantly extends Giotto Suite’s capabilities for spatial downstream analysis and visualization.

## Discussion

We present a new generation spatial analysis framework, Giotto Suite, which offers a fully integrated and comprehensive suite of tools that were built to provide end-to-end workflows encompassing every critical stage of working with data generated by the latest spatial technologies. In this manner, Giotto Suite differs significantly from the original Giotto package^21^ and provides an all-in-one solution that is otherwise only offered through a combination of multiple recently developed tools^56–61^.

First, Giotto Suite adheres to its technology-agnostic approach by providing data ingestion pipelines that are compatible with any type of raw data structure. At the core of the Giotto Suite framework, we developed innovative data classes to represent biological datasets across multiple spatial and data modalities. In addition, these data classes form their own fully independent sub-objects and can thus be the starting point for independent spatial workflows, such as providing solutions for imaging-based segmentation-free clustering approaches^62^. Importantly, these core classes make it possible to generate a multi-modal hierarchical representation that faithfully reflects the organizational principles of tissue architecture and provides a unified approach for data representation. Similarly, it underlies our seamless integration with co-registration methods such that multiple spatial technologies can be jointly queried or analyzed together. In this manner, our approach provides more flexibility than the recently developed *SpatialData*^58^ package, which enforces a standard data framework. Notably, to accommodate the increasing size of spatial multi-modal datasets, we developed *GiottoDB*, which provides the groundwork that developers and users can use to represent their data through different backends that can scale according to their needs.

Next, Giotto Suite provides users with a modular set of tools for spatial analysis and visualization, including a responsive coding environment and seamless interoperability to both spatial and other genomics software communities. In this manner, Giotto Suite provides direct access to other spatial omics ecosystems such as those from R/Bioconductor^52,57,60^ and geospatial^55^ communities. Although similar effort has been taken in *Voyager*^57^, the underlying `SpatialFeatureExperiment` class currently limits working with multiple spatial layers at different scales or with different data modalities. Hence, Giotto Suite provides a more integrated and inclusive solution that combines the strengths of these methods.

Altogether, Giotto Suite offers an ideal benchmarking framework to systematically implement and compare a large number of new spatially related methods. The results of such analyses are needed for establishing best practices and data standards. It is also well suited to adopt and adhere to any future minimum information guidelines^63^ or metadata standards^64^ that will be necessary for spatial dataset and method harmonization. These activities are especially important for coordinating large-scale collaborative efforts that involve many technology and data analysis experts, such as in various cell atlas projects^6,48,65,66^.

Finally, we have demonstrated the utility of Giotto Suite through applications to multiple spatial datasets from a diverse set of technologies that span across various spatial resolutions and modalities. Taken together, the core framework and new tools implemented in Giotto Suite provide a powerful solution for a seamless harmonization and integration of diverse datasets.

## Contributions

J.G.C., J.C., G-C. Y. and R.D. conceived the project. J.G.C., J.C., M.O.B., J.X., E. R., W.W., I. A., I.S., P.G., A.S., and R.D. developed Giotto Suite, implemented and revised code. J.G.C., J.C., M.O.B., J.X., E. R., W.W. performed analyses and generated figures. G-C.Y. and R.D. wrote the manuscript with input and feedback from all other authors.

## Data and Code Availability

The packages that are part of Giotto Suite can be downloaded from our GitHub page http://github.com/drieslab and additional information, including a large set of examples, vignettes, and FAQs can be found on http://giottosuite.com.

The following spatial datasets were used in this manuscript:

*Spatial Genomics dataset.* The mouse kidney fresh frozen dataset was downloaded from the Spatial Genomics website at https://spatialgenomics.com/data/.

*DBiT-seq dataset*. The mouse embryo E10.5 dataset was downloaded from https://www.ncbi.nlm.nih.gov/geo/query/acc.cgi?acc=GSE137986.

*Nanostring CosMx dataset.* The CosMx FFPE Non-Small Cell Lung Cancer dataset for lung sample 12 was downloaded from the Nanostring website at https://nanostring.com/products/cosmx-spatial-molecular-molecular-imager/ffpe-dataset/nsclc-ffpe-dataset/

*Seq-Scope dataset*. The Seq-Scope liver dataset was downloaded directly from the following link, https://deepblue.lib.umich.edu/data/concern/data_sets/9c67wn05f.

*Vizgen dataset.* The MERSCOPE/MERFISH FF mouse brain (data release v1.0, May 2021) and FFPE breast cancer (May 2022) datasets were downloaded directly from the Vizgen website at https://info.vizgen.com/mouse-brain-data and https://info.vizgen.com/ffpe-showcase, respectively.

*10X Genomics Xenium dataset.* Xenium, corresponding images and Visium datasets for human breast cancer were downloaded directly from the 10X website at https://www.10xgenomics.com/products/xenium-in-situ/preview-dataset-human-breast.

*10X Genomics multi-modal Visium CytAssist Human Tonsil dataset.* The multi-modal Visium CytAssist Human Tonsil dataset was downloaded from the 10X genomics website at https://www.10xgenomics.com/resources/datasets/gene-protein-expression-library-of-human-tonsil-cytassist-ffpe-2-standard.

*10X Genomics Visium Mouse Brain Section (Coronal) dataset.* The Visium Mouse Brain Section (Coronal) dataset was downloaded from the 10X Genomics website at https://support.10xgenomics.com/spatial-gene-expression/datasets/1.1.0/V1_Adult_Mouse_Brain

*Stereo-seq dataset.* The bin1 matrix files (i.e. *_GEM_bin1.tsv.gz) were downloaded from the CNGB portal at the following link (https://db.cngb.org/stomics/mosta/download/).

*Imaging Mass Cytometry dataset*. Intensity images of human lymph node FFPE tissue were downloaded from a repository created by Bost et. al https://data.mendeley.com/datasets/ncfgz5xxyb/1.

*Single Cell Mouse Brain Dataset.* A single-cell reference dataset published by <u>Manno et al. 2021</u> was used to identify developmental mouse brain cell types for spatial DWLS deconvolution with Stereo-seq data in this study. This data can be downloaded in the form of a .loom file from the Mousebrain.org website (http://mousebrain.org/development/downloads.html).

*Single-cell Human Tonsil Dataset*. The Atlas of Cells in the Human Tonsil published by <u>Massoni-Badosa</u> <u>et al 2022</u> containing the annotation of >357,000 cells was used for spatial DWLS deconvolution of Visum CytAssist data in this study. The annotated SpatialExperiment object was downloaded from https://github.com/massonix/HCATonsilData.

## Acknowledgments

We would like to thank former and current members of the Dries and Yuan labs for their support, contributions, and invaluable feedback on the Giotto Suite project and development over time. More specifically we would like to thank the following individuals: Emma Kelley, Christina Ennis, Jason Weis, Yibing Michelle Wei, Rafael dos Santos Peixoto, Mohammed Muzamil Khan, Jason Yeung, Amelia Zug, Laila Norford, Cecilia McCormick, Natalie Del Rossi, Azra Krek, Weiping Ma, Xuan Cao, and Shiwei Zheng.

## Supplementary Figures

**Supplementary Figure 1:**
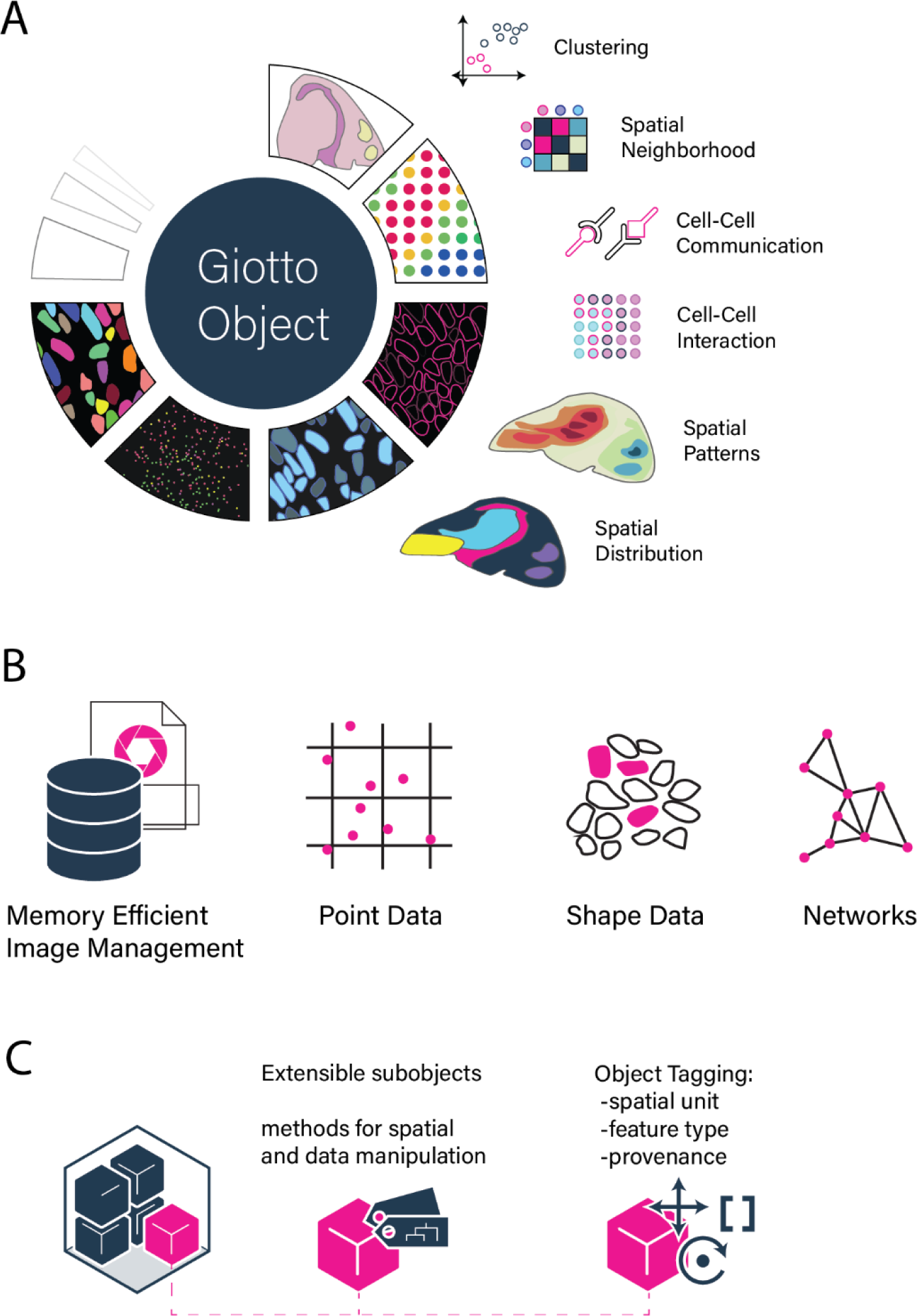
Giotto Suite workflows and data structures. **A)**An overview of the types of spatial analyses and workflows implemented in Giotto Suite. **B)** Pictograms depicting the various Giotto Suite data representations within a typical spatial data analysis pipeline. **C)** Schematic representation of sub-object creation from defined feature types and spatial units with encoded provenance information.

**Supplementary Figure 2:**
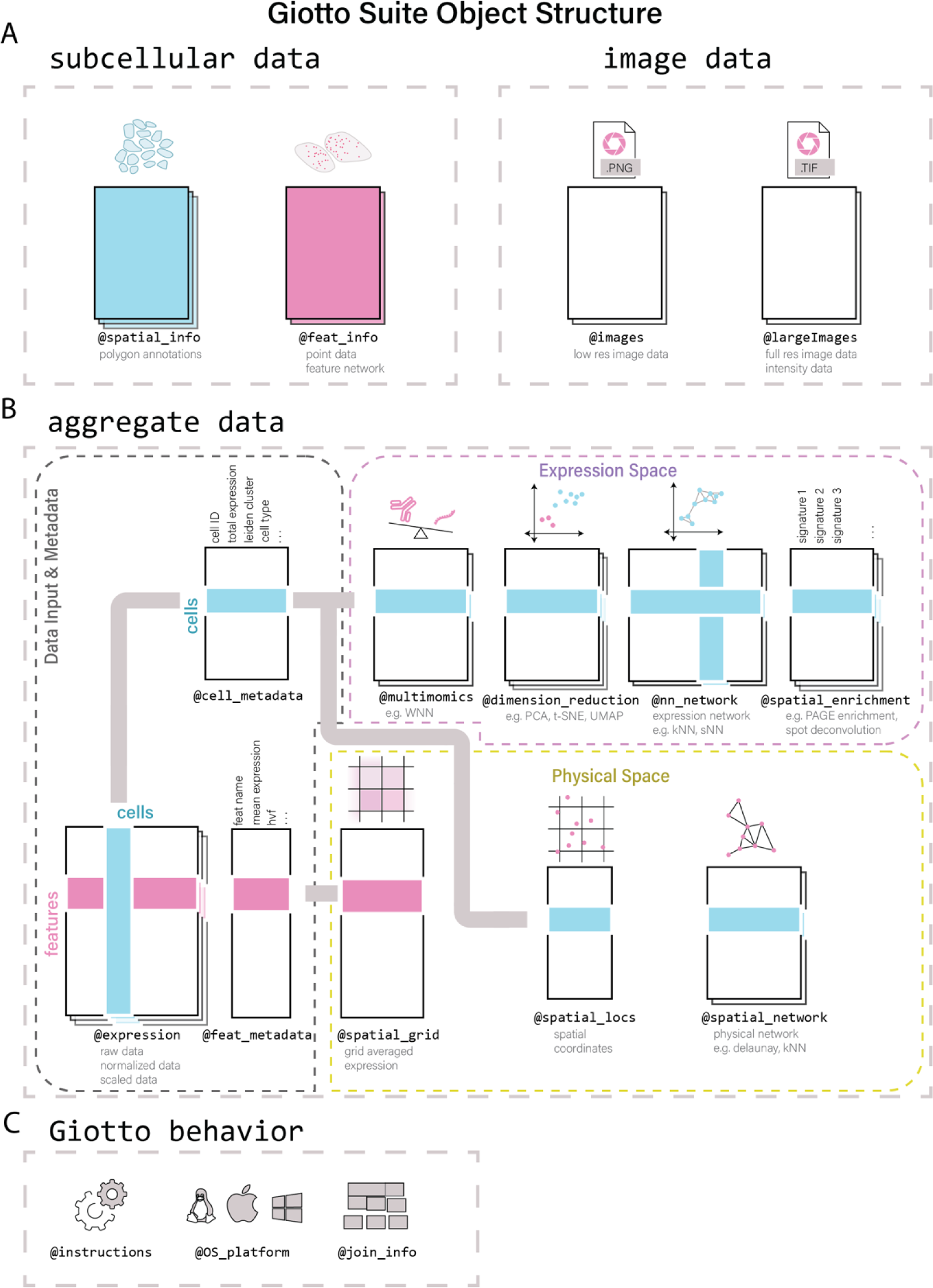
Giotto Suite subcellular data organization and aggregation. **A**) Schematic depicting the storage of different subcellular and image data types. **B**) Visualization of Giotto Suite’s object slots that are available to store different raw and processed information from aggregated spatial data. Aggregate information can be created from any spatial unit and feature type information encoded in the Giotto object spatial_info and feat_info slots in A, respectively. **C**) Additional tools available in Giotto Suite to dictate default behavior and functionalities in downstream spatial analysis workflows.

**Supplementary Figure 3:**
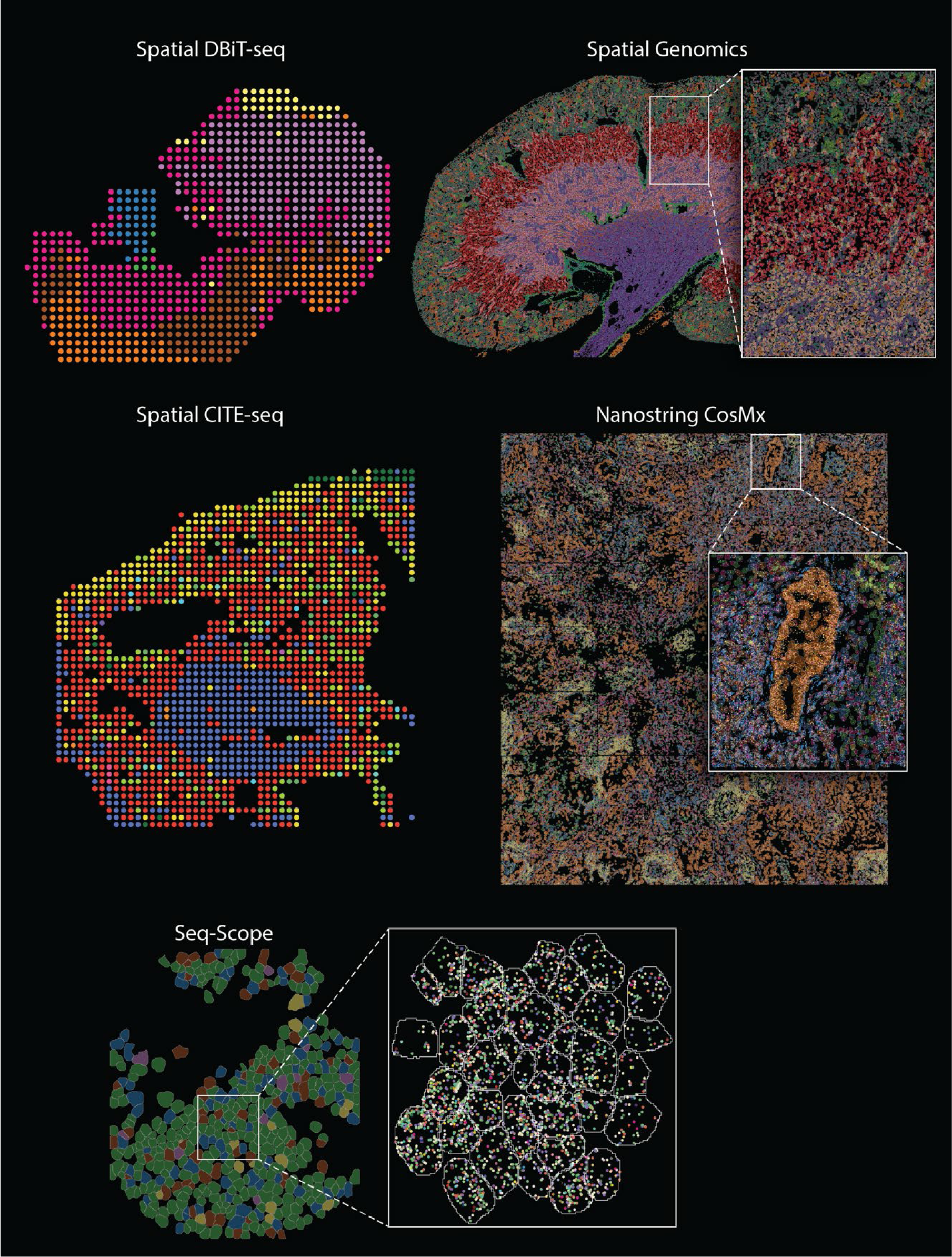
Technology-agnostic identification of spatial expression patterns. Identification and visualization of expression patterns identified through spatial co-expression analysis in datasets from various recently developed spatial technologies. Zoomed-in regions highlight transcript information at the subcellular level.

**Supplementary Figure 4:**
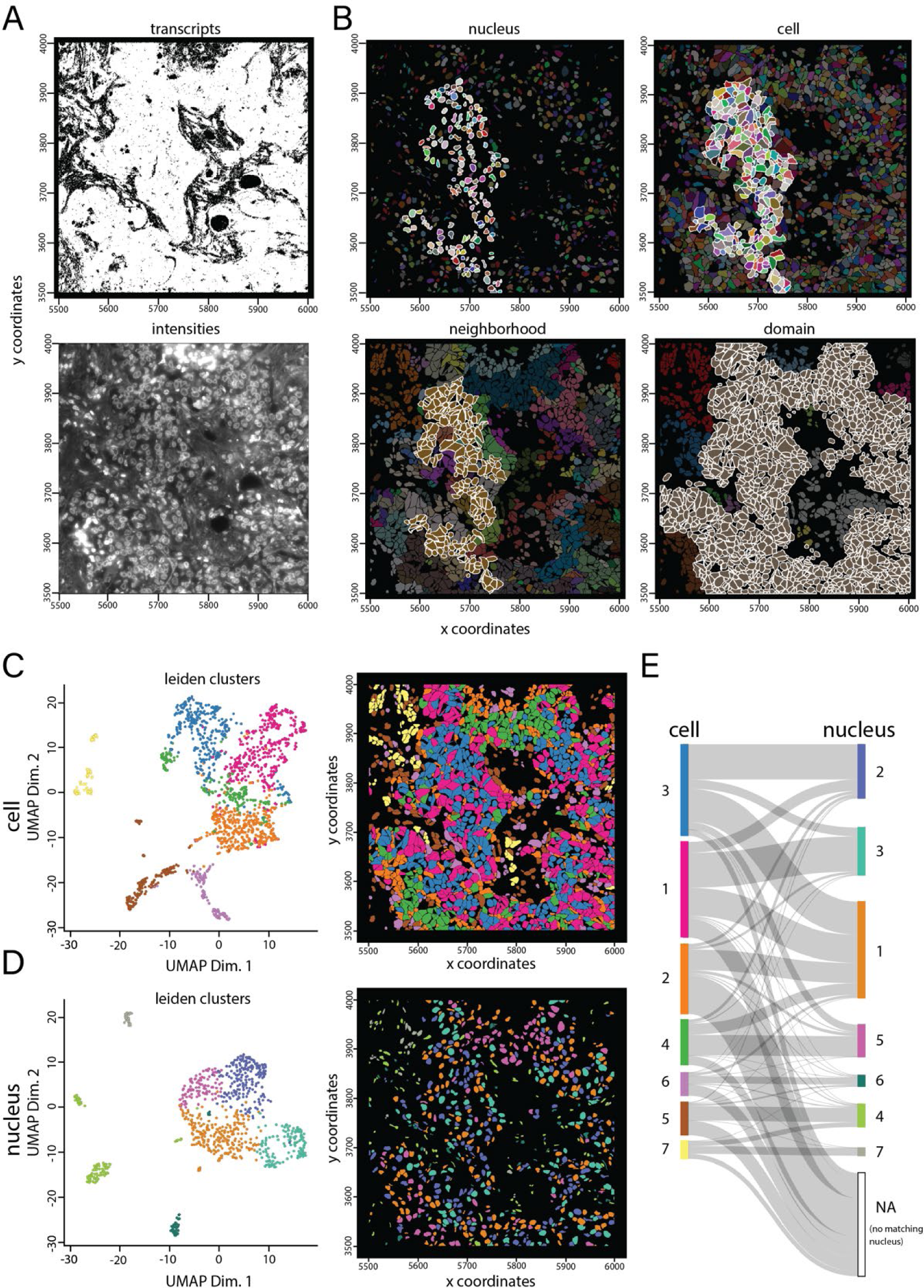
Multi-scale analysis. **A)**Spatial plots showing raw transcript locations (top) and corresponding DAPI image (bottom). **B)** Spatial plots illustrating annotations at different scales, including nucleus (top left), cell (top right), neighborhood (bottom left), and domain (bottom right). Highlighted annotations correspond with selection in Fig. 2B. C-D) UMAP and spatial plot showing Leiden clusters for cellular (**C**) and nuclear (**D**) segmentations. **E)** Sankey plot showing the Leiden cluster relationships between count aggregation results from cellular (**C**) and nuclear (**D**) segmentations.

**Supplementary Figure 5:**
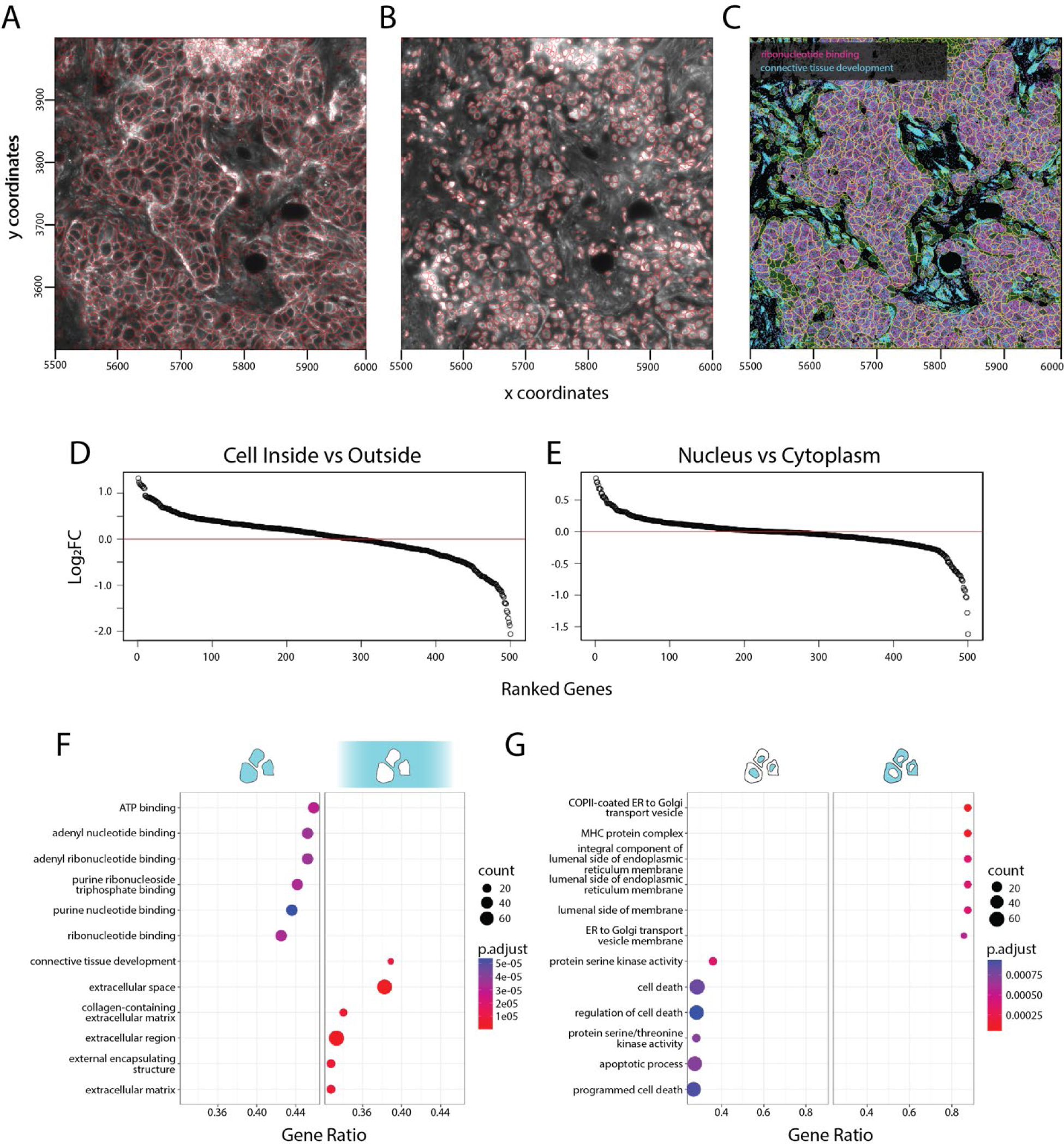
Subcellular gene set enrichment analysis. **A-B)** Spatial plots showing cellular (**A**) and nuclear (**B**) segmentation results in red. **C**) MERFISH dataset subset depicting cellular (magenta) and extracellular (cyan) enriched transcripts for all genes in GSEA terms related to “ribonucleotide binding” and “connective tissue development”, respectively. **D-E**) Waterfall plot depicting the log-fold enrichment changes for all transcripts within versus outside of the cell (**D**) and nuclear versus cytoplasm (**E**). **F-G**) GSEA enrichment plots showing results for inside versus outside cells (**F**) and nuclear versus cytoplasm (**G**).

**Supplementary Figure 6:**
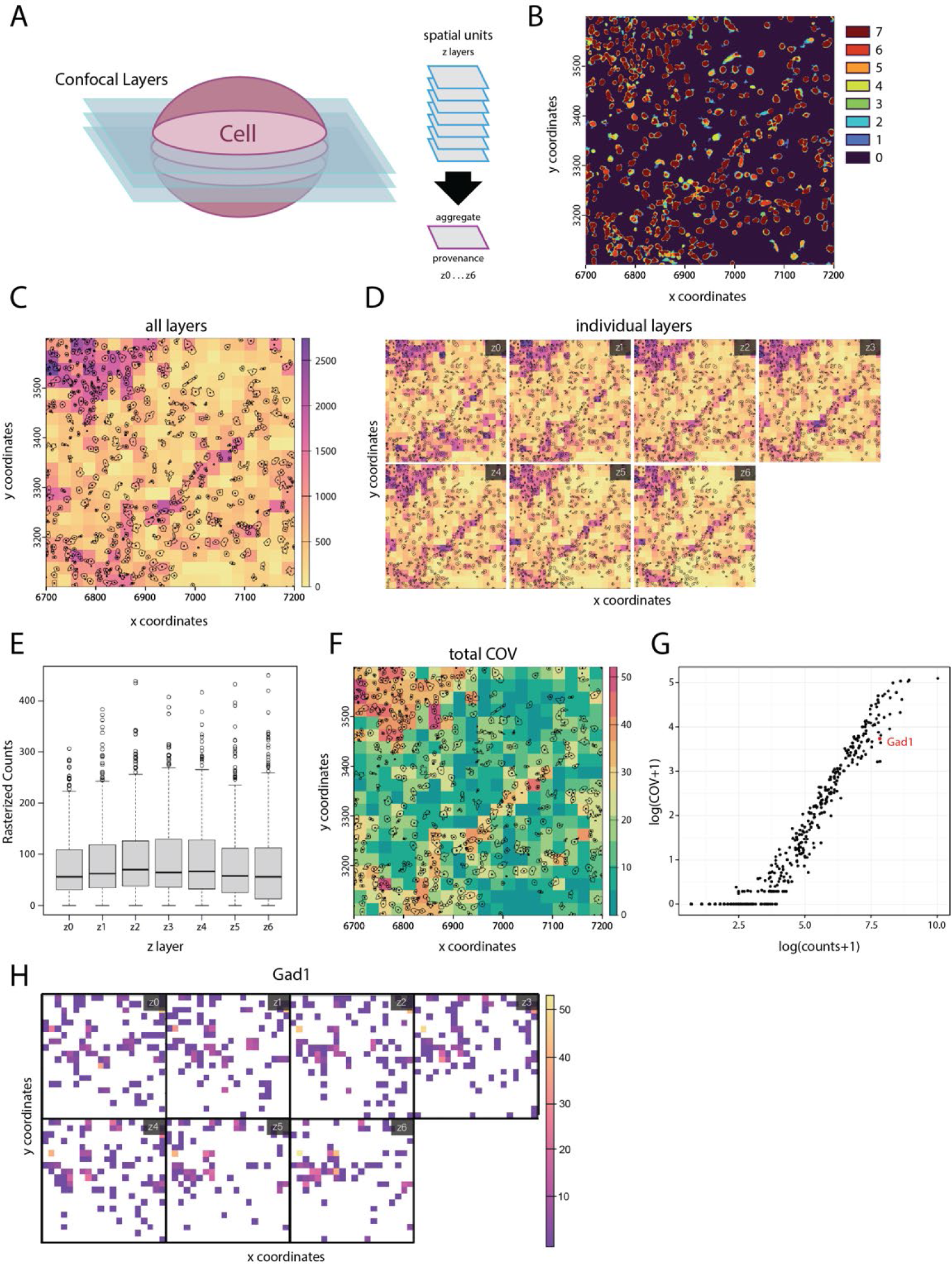
Transcript detection variance across z stacks. **A)** Pictogram of z stacks each with their own detected transcripts and cellular segmentation information. **B)** Spatial plot depicting the overlapping polygons from all z stacks combined. Colors represent how many polygons overlap at each pixel. **C)** Summed and rasterized expression combined for all layers and transcripts. **D)** Summed and rasterized expression for all transcripts within each individual layer. **E)** Boxplots showing summed rasterized expression levels for all pixels within each layer. **F)** Coefficient of variation (COV) for expression levels from all transcripts per rasterized pixel across all layers. **G)** Scatterplot showing log counts versus log COV across all z stack layers for each gene. **H)** Rasterized expression levels per pixel for Gad1 for each z stack layer.

**Supplementary Figure 7:**
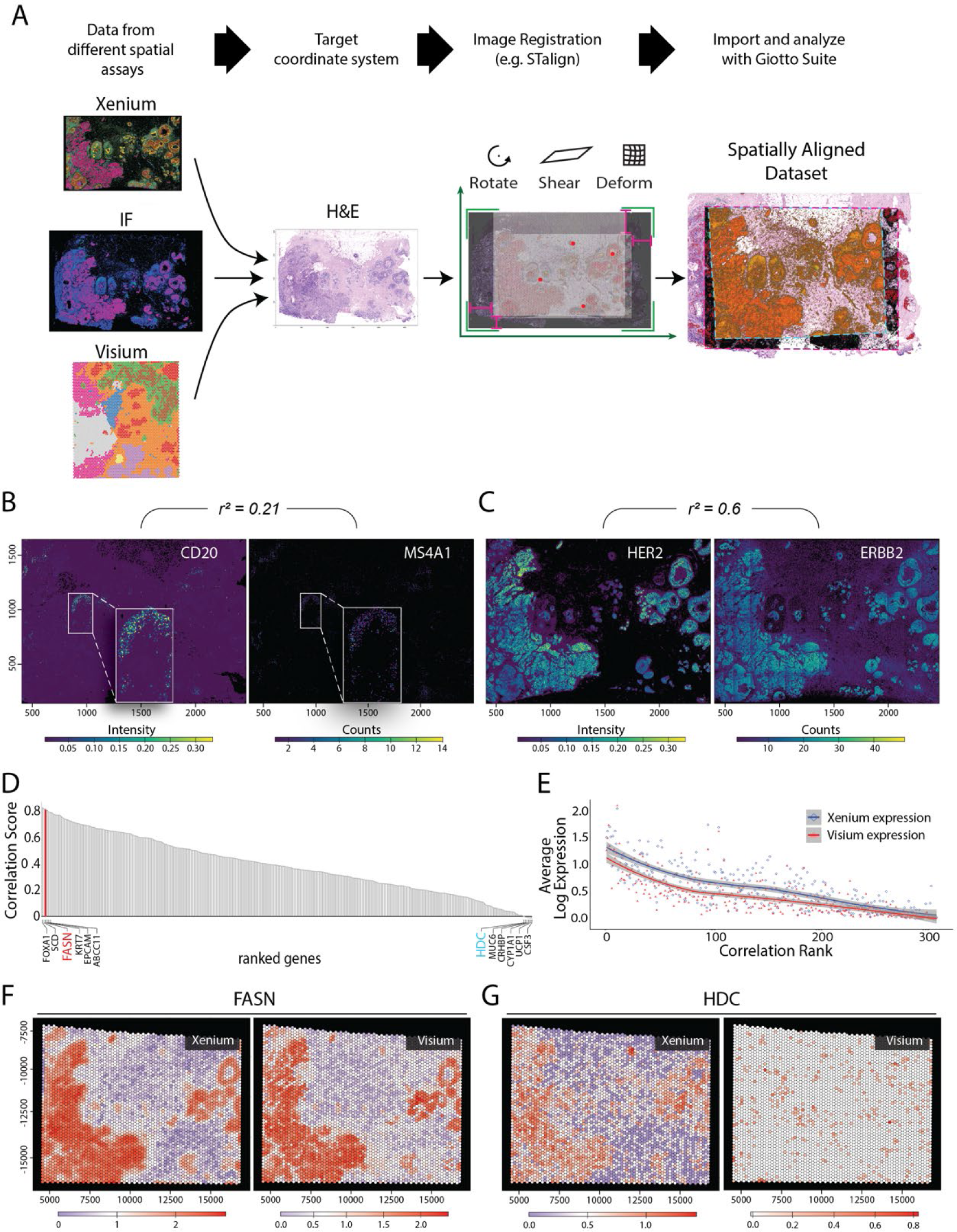
Comparison of co-registered multimodal datasets. **A**) A schematic workflow of the co-registration step for the 10X Genomics Xenium pre-release breast cancer dataset. **B**) Rasterized CD20 intensity data (left) and rasterized MS4A1 transcript counts (right). Pearson’s r is shown. **C**) Rasterized HER2 intensity data (left) and rasterized ERBB2 transcript counts (right). Pearson’s r is shown. Rank of correlation scores between overlapping genes expressed in Visium and registered Xenium data. **D)** LOESS regression (solid curves) of log expression against ranked correlation scores from **(D)**. **F-G**) Spatial feature expression plots for registered Xenium and Visium data for genes FASN (**F**) and HDC (**G**).

**Supplementary Figure 8:**
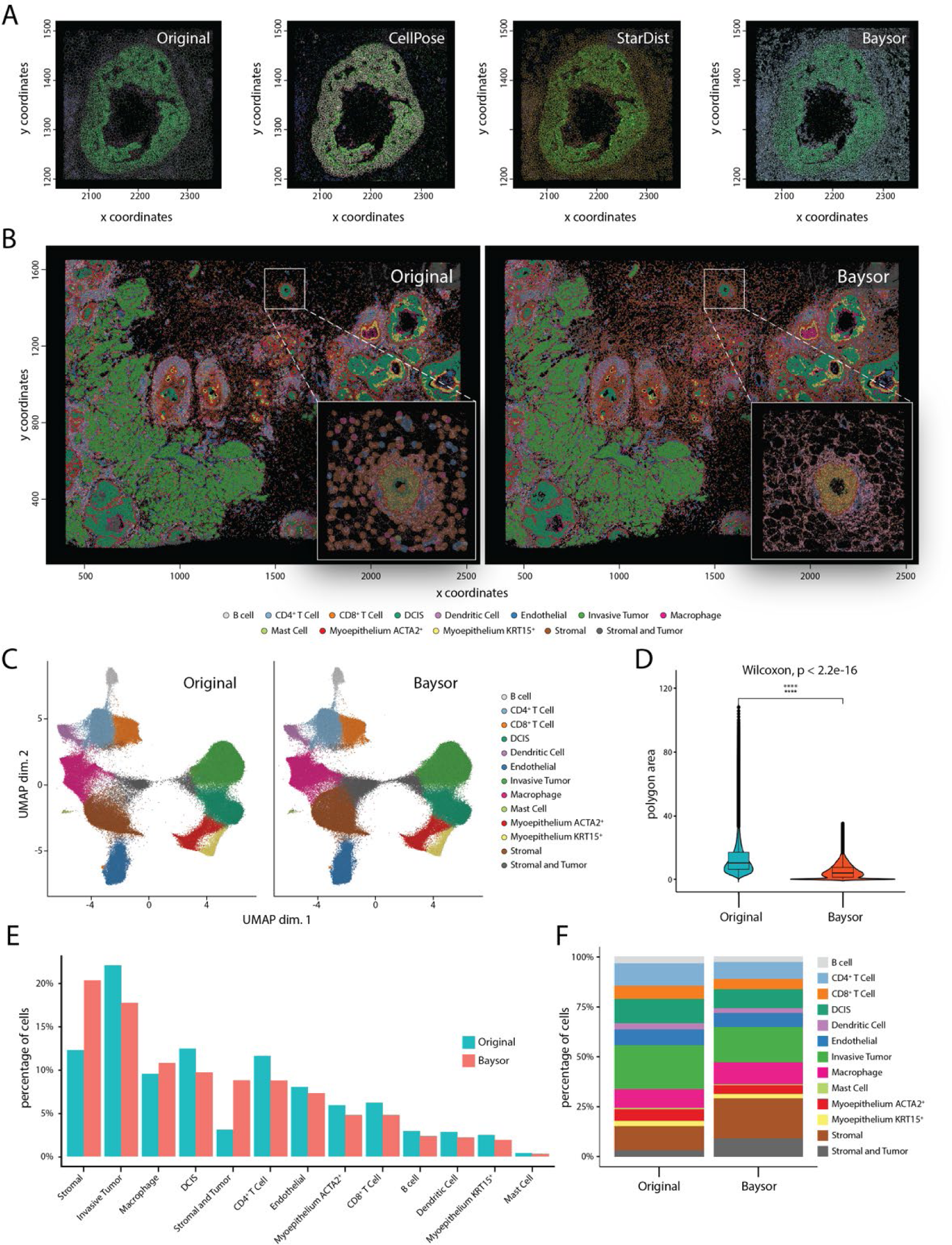
Joint analysis of multiple cellular segmentation results. **A)** Spatial plots depicting the segmentation results from a set of established segmentation methods on a subset of the 10X Genomics Xenium Breast Cancer data. **B)** Spatial plots of the 10X Genomics Xenium Breast Cancer data depicting the segmentation results and cellular annotations for the original (left) and Baysor (right) segmentation methods. **C)** UMAP plots showing the joint clustering results for both the manufacturer-provided and Baysor cellular segmentations. **D)** Violin plots showing the surface area of polygons identified by the original and Baysor segmentation methods. **E)** Barplot illustrating the percentage of cellular annotations for both the original and Baysor segmentation results. **F)** Stacked barplot showing the percentage of cellular annotations for both the original and Baysor segmentation results.

**Supplementary Figure 9:**
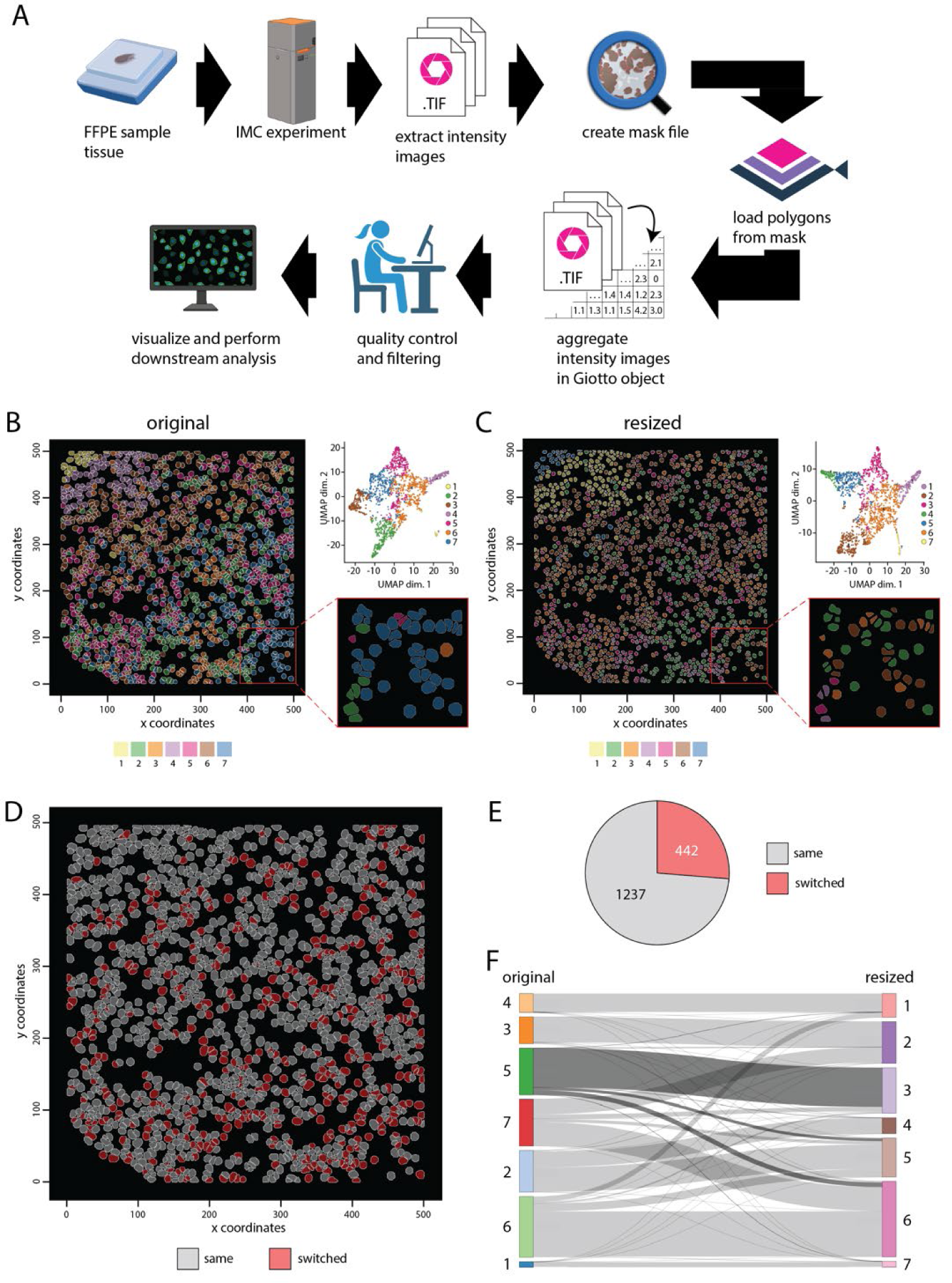
Polygon resizing effect on cell annotations. **A)**Example of a multi-step end-to-end workflow for processing, analyzing, and visualizing imaging mass cytometry (IMC) data. **B**) Spatial and UMAP plots showing the cell clustering and annotation results from kmeans (k = 7). **C**) Spatial and UMAP plots showing the cell clustering and annotation results from kmeans (k = 7) starting from 25% downscaled cell polygons compared to **(B)**. **D**) Spatial plot depicting cell IDs that stayed in the same (gray) or changed (red) their majority cluster annotation. **E**) Pie chart showing the number of changed cell annotations between the original and downscaled cell polygons. **F**) Sankey diagram illustrating the relationship between cell annotations from the original versus downscaled cellular polygons.

**Supplementary Figure 10:**
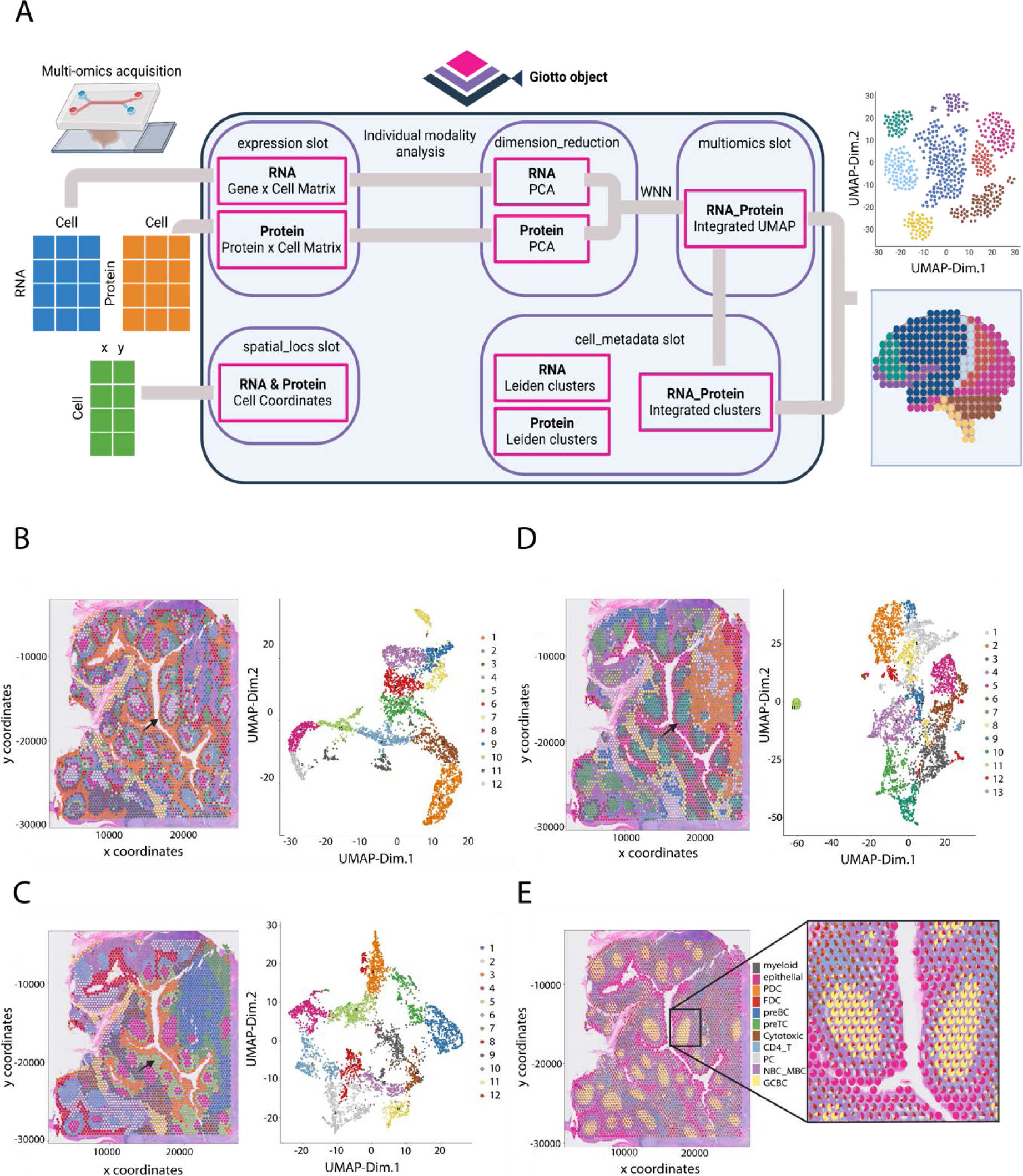
Multi-omics data analysis using the Giotto Suite framework. **A**) Schematic representation and workflow for multi-modality integration of RNA and Protein spatial transcriptomic datasets. **B-E**) 10X genomics Human tonsil dataset. Analysis at the RNA (**B**), Protein (**C**) level, multi-omics integration (**D**), and DWLS deconvolution (**E**) using a reference scRNAseq were performed.

**Supplementary Figure 11:**
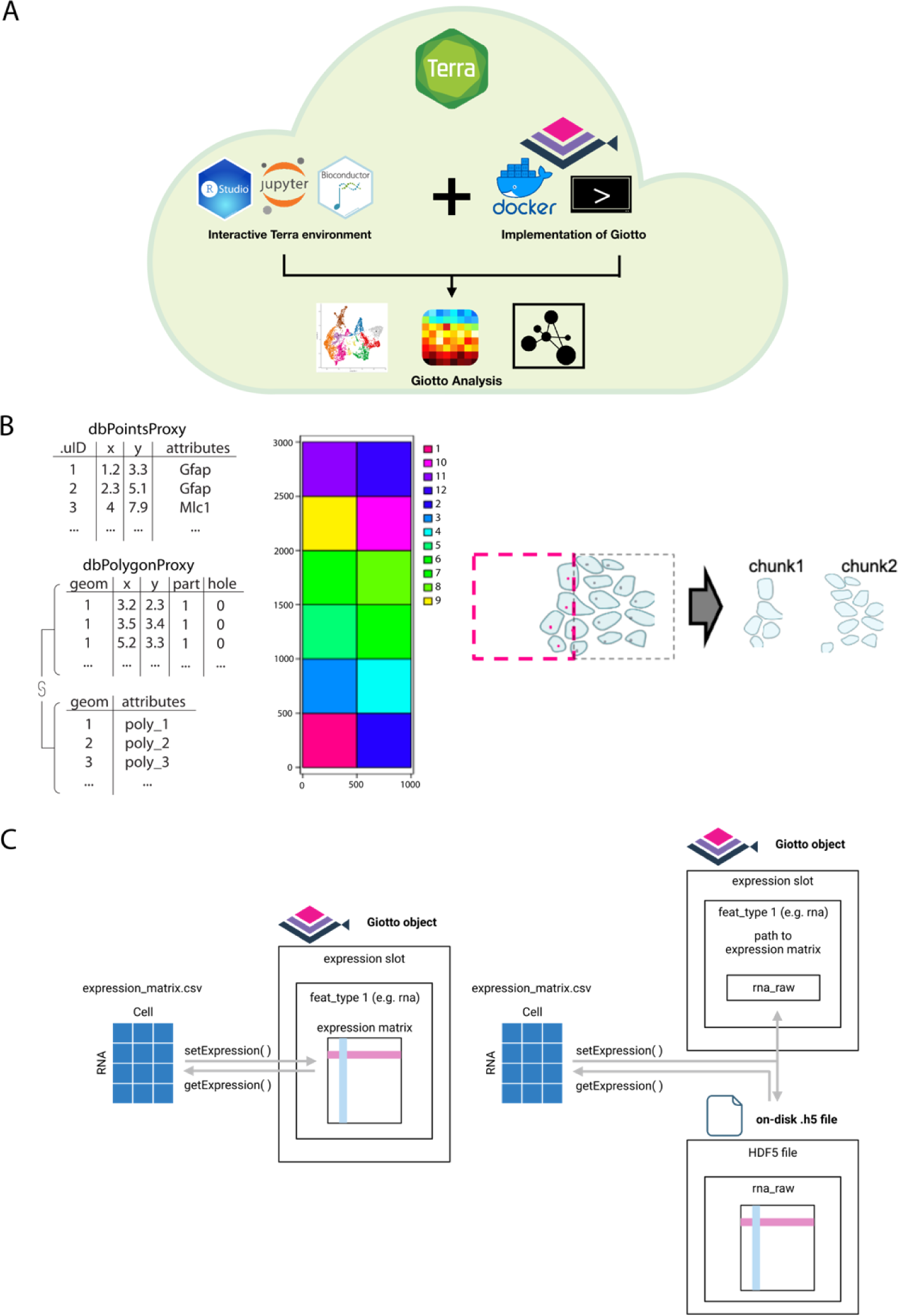
Scalability implementations for Giotto Suite. **A**) Docker images and startup scripts compatible with terra.bio platform were developed to facilitate running analysis using Giotto on the cloud. The implementation creates a customized cloud environment ready to run Giotto analysis using interactive Jupyter notebooks and the RStudio app. **B**) Pictogram showing the implemented chunking strategy utilized by Giotto Suite to perform large spatial analytical operations. **C**) Schematic for the implementation of the delayed HDF5 backend. The standard sparse expression matrix (left) is replaced within the ‘expression’ slot with a string that indicates the internal path to the expression matrix within an on-disk .h5 file (right).

**Supplementary Figure 12:**
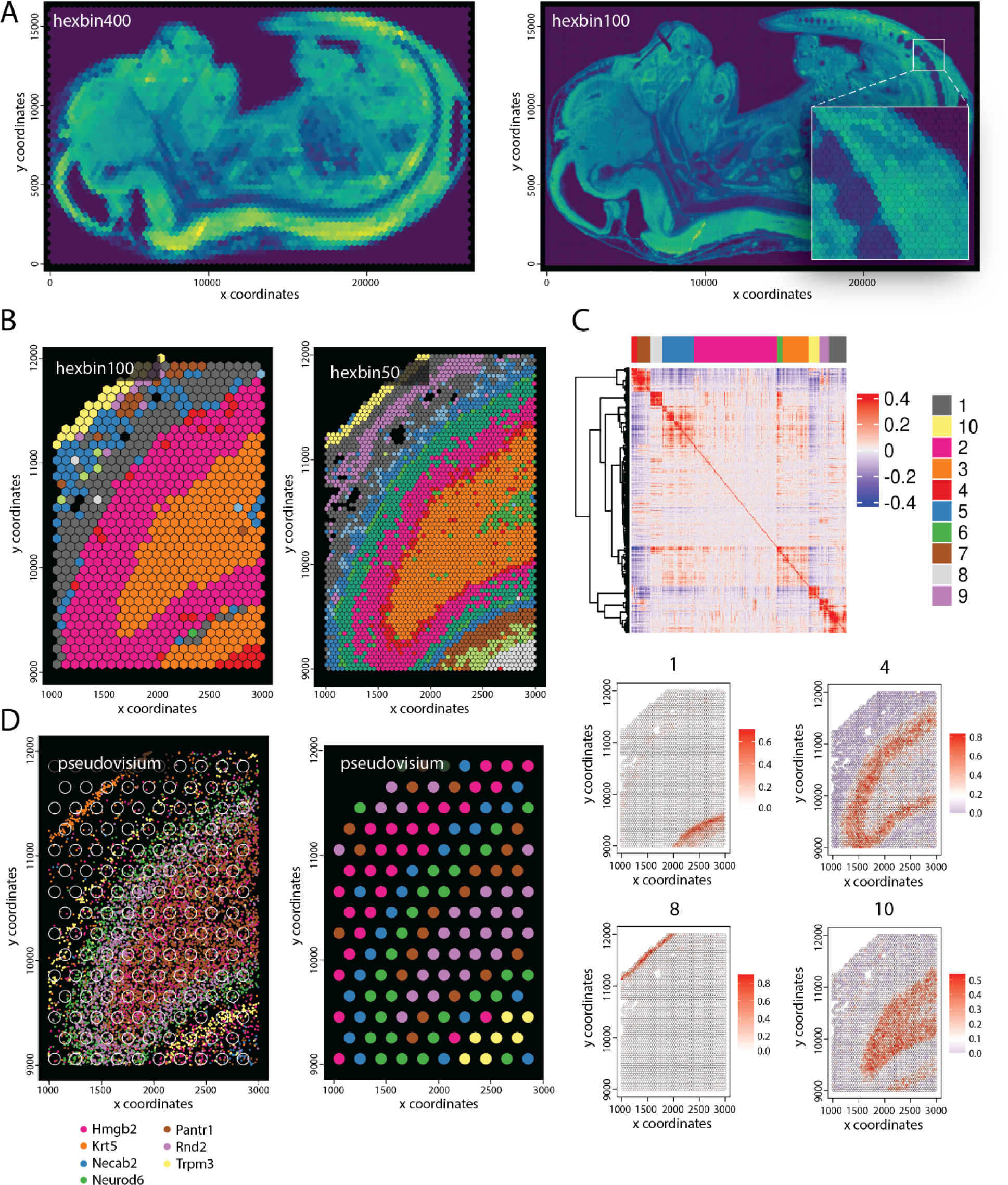
Tiling approaches and applications. **A)**Spatial plots showing aggregated stereo-seq transcript counts for hexagon tiles with diameter = 400 (left) and 100 (right). Inset shows zoomed-in tail region. **B)** Spatial plots showing Leiden clustering results at low (hexagon, diameter = 100) and high (hexagon, diameter = 50) tile resolution for the selected stereo-seq region in Fig. 2J. **C)** Heatmap (top) depicting spatial gene co-expression modules for the selected region in B. Spatial metagene plots (bottom) showing the expression of selected spatial co-expression modules. **D)** Visualization illustrating the creation of a Pseudo-Visium dataset from a stereo-seq dataset. Spots with diameter 55 µm and inter centroid distance equal to 100 µm (left) were used to aggregate individual transcripts at bin1 level. This was followed by downstream spatial processing steps and the generation of Leiden clusters at the spot level (right).

**Supplementary Figure 13:**
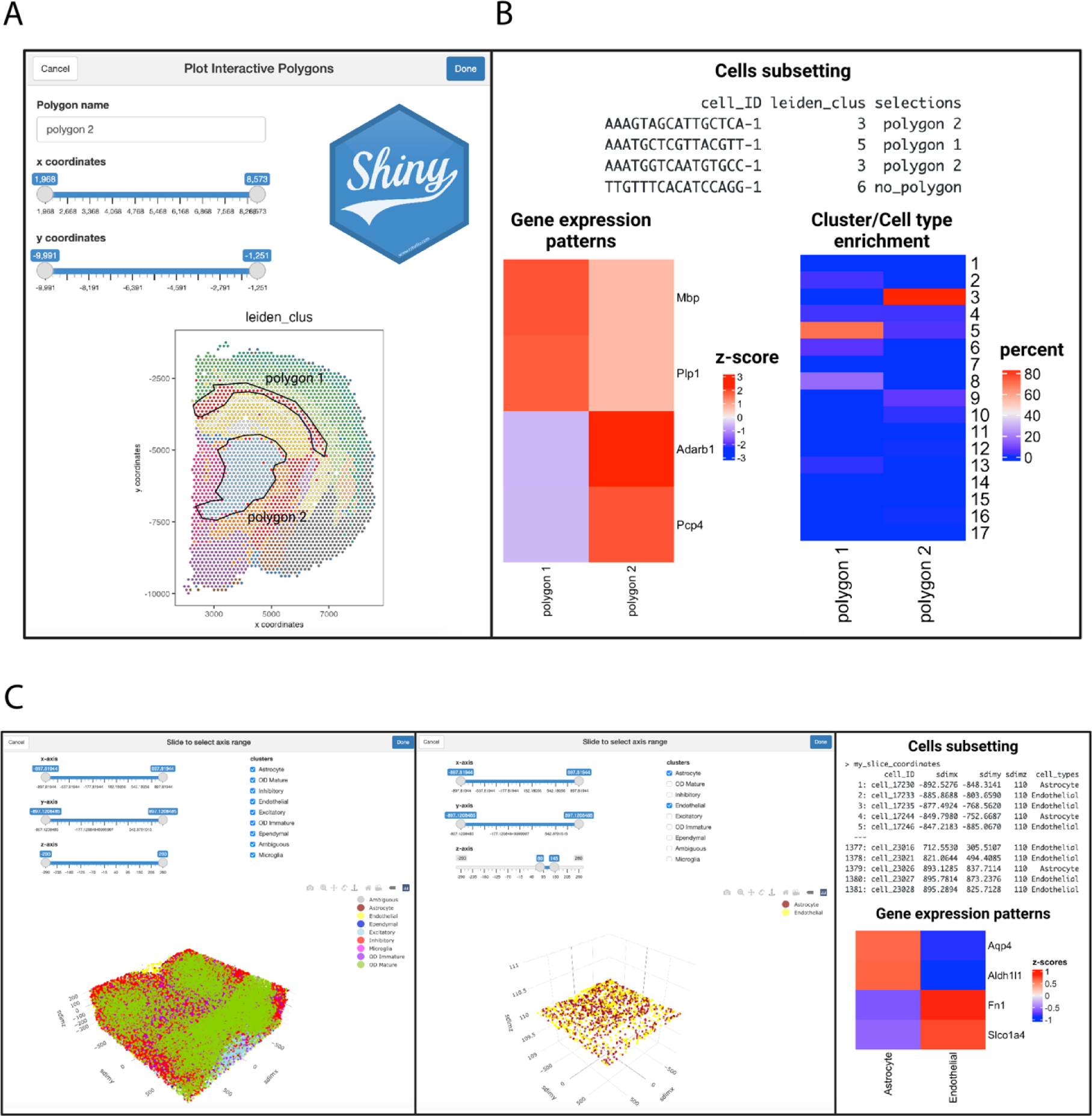
Interactive R/Shiny applications facilitate the selection and comparison of regions of interest. **A**) A local-running Shiny gadget was developed to interactively select regions of interest over a Giotto Suite spatial plot. **B**) The information from plotted regions of interest is used for downstream analysis such as subsetting of cells, as well as gene and cell type enrichment comparison. **C**) An interactive tool for plotting and subsetting tri-dimensional datasets was developed. The tool facilitates subsetting across the z-axis to create slices and select cell types. Downstream analyses include Giotto object subsetting and feature enrichment.

**Supplementary Figure 14:**
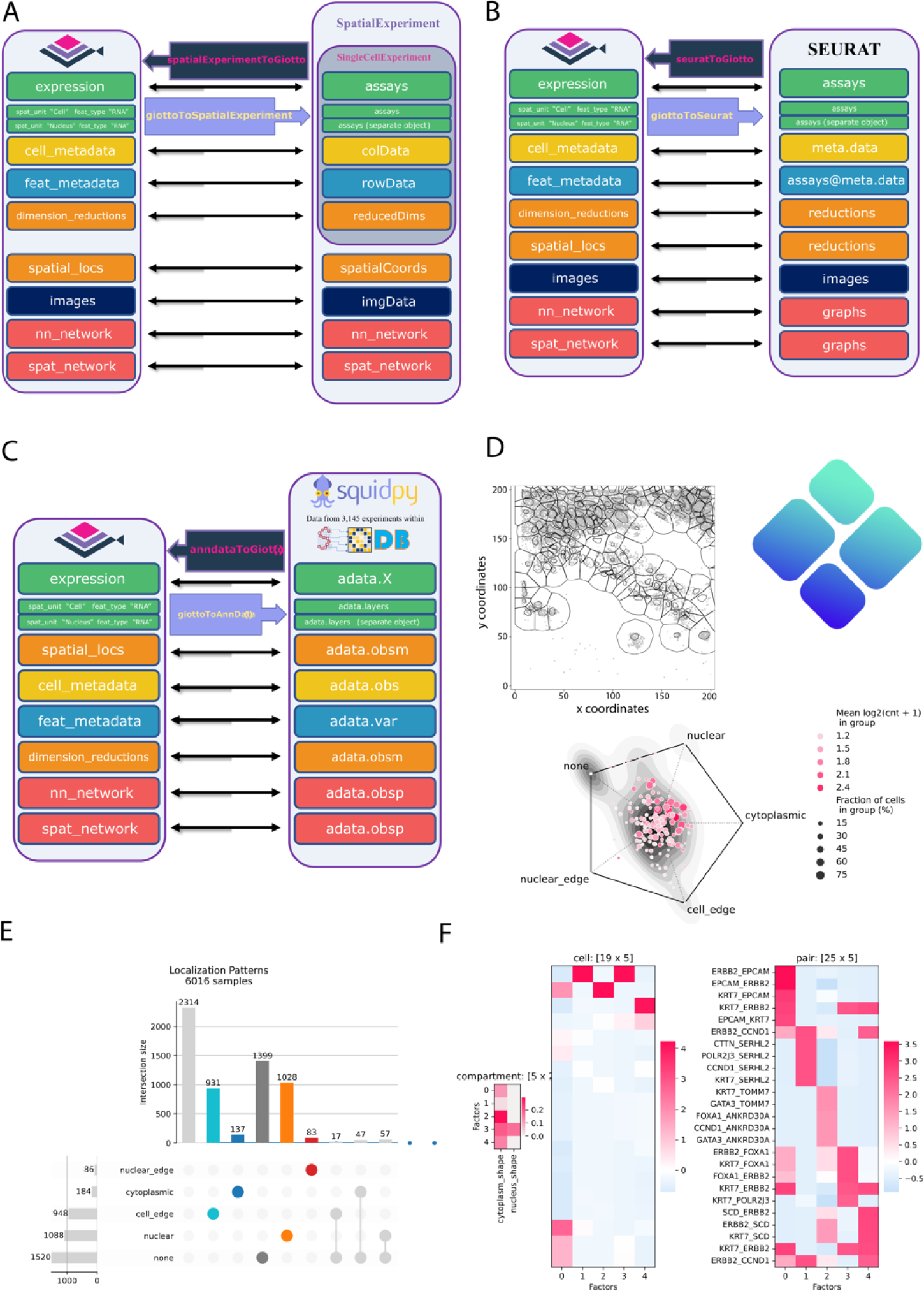
Spatial data analysis community interoperability. **A-C**) Schematics depicting the mapping strategies between a Giotto object and a **(A)** SpatialExperiment, **(B)** Seurat, or **(C)** AnnData/Squidpy object. **D**) The spatial plot (top) shows the region selected for Bento analysis. RadViz plot (bottom) depicts the spatial subcellular distribution for all transcripts. **E**) UpSet plot showing the number of transcripts with different spatial distribution features. **F**) Heat maps showing the results of Bento colocalization analysis. In this analysis, 5 colocalization factors were identified. Heatmaps showing the loading of each colocalization factor on spatial features (left panel), cell distributions (middle panel), and gene pairs (right panel).

**Supplementary Figure 15:**
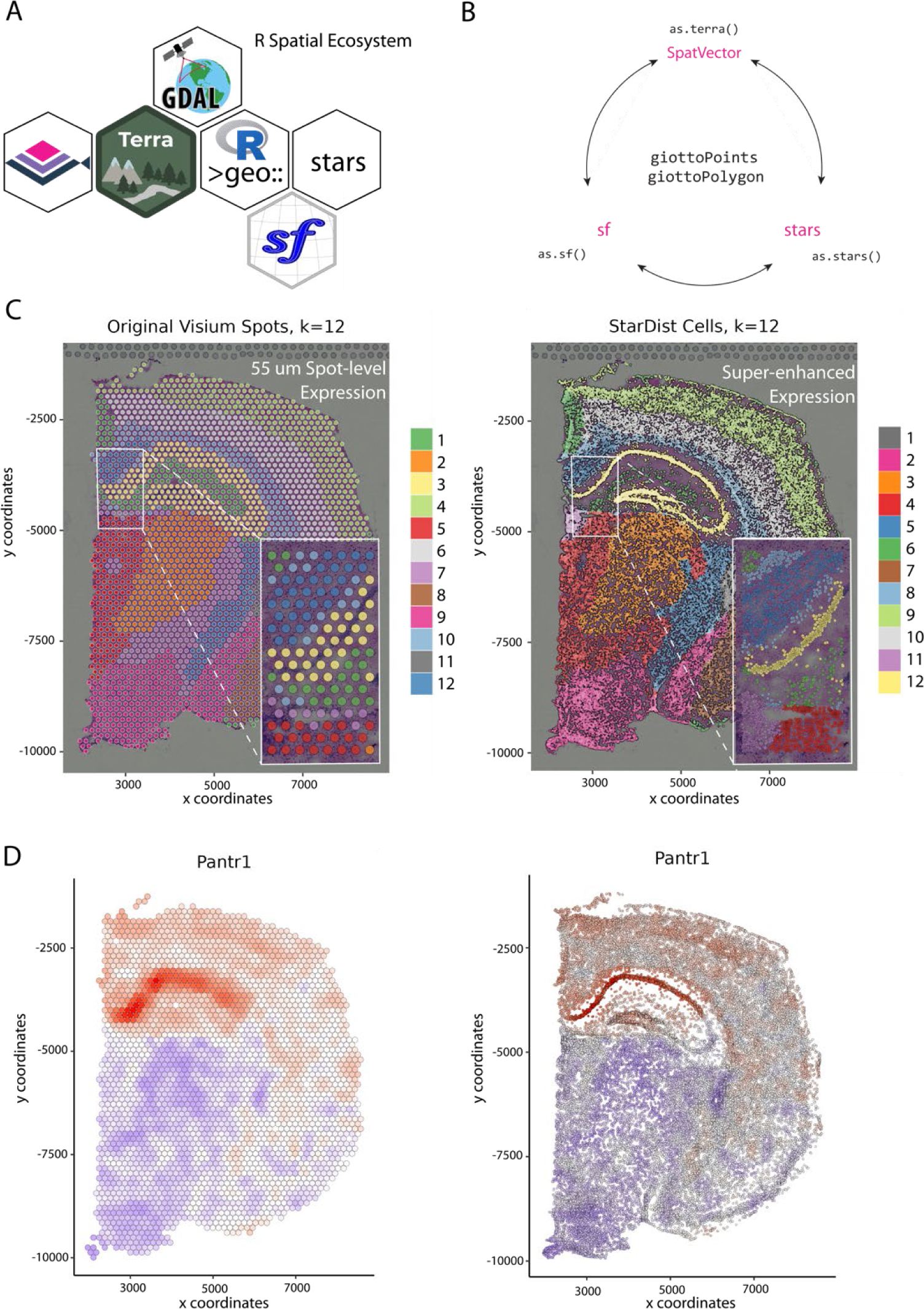
Integration of Giotto Suit with the R (geo)spatial ecosystem. **A)**Pictogram illustrating the connection between Giotto Suite and the larger R geospatial ecosystem. **B**) Giotto-native ‘as’ functions for converting giottoPoints and giottoPolygon to corresponding *terra*, *sf*, and *stars* classes **C**) Spatial plots showing spatial clustering results from original (left) and super-enhanced (right) Visium gene expression data. Insets depict zoomed-in regions illustrating differences in resolution (55 µm vs single-cell). **D**) Spatial gene expression plot for Pantr1 from original (left) and super-enhanced (right) Visium gene expression data.

## Methods

### Spatial data co-expression processing pipeline

A standard Giotto spatial data processing and analysis workflow was used to visualize spatial clusters starting from a balanced set of genes from spatial co-expression modules. A similar pipeline was used for data generated by Nanostring CosMx, Spatial Genomics, DBiT-seq, Spatial CITE-seq, and Seq-Scope. More specifically, raw data was loaded followed by filtering, normalization, and detection of spatial genes using `binSpect()`. Top spatial genes (`max = 500`) were subsequently used to create spatial gene co-expression modules followed by hierarchical clustering and the selection of an equal representation of spatial gene groups from each co-expression module with `getBalancedSpatCoexpressionFeats()`. These genes were then used in a typical Giotto Suite pipeline, including dimension reduction (PCA), creation of a shared nearest-neighbor network, and Leiden clustering to create spatial expression-informed clusters for each tissue sample.

### MERSCOPE data processing and analysis

Two Vizgen MERSCOPE datasets were used. The FFPE Human Immuno-oncology Breast Cancer (patient 1) dataset and the Mouse Brain Receptor Map (v1.0. May 2021 Slice 1, replicate 1). A key difference between both datasets is that the Breast Cancer dataset contains cellular segmentations, while the Brain dataset contains nuclear segmentations. In addition, the Brain dataset also provides different segmentation results associated with each of the 7 provided z stacks.

For each dataset, associated images were first loaded in as `giottoLargeImages` and mapped to microns using the provided “transforms” information via `createMerscopeLargeImage()`, then cell segmentations and transcript detections were aligned to the same coordinate reference frame. These early MERSCOPE datasets provide cell segmentation information as a directory of thousands of HDF5 files, separated by FOVs. The files were first scanned through to produce an `H5TileProxy` object that contains a spatially indexed manifest of the files, their contents, and a parse function for converting chunks of data read in from the HDF5 files into the format expected downstream. This provided a framework for spatially chunked access to the polygon information. For visualization purposes and faster processing of the analysis examples the datasets were spatially subset to 500 x 500 micron regions. The breast cancer dataset was subset to 5500, 6000, 3500, 4000, and the mouse brain dataset was subset to 6700, 7200, 3100, 3600 (xmin, xmax, ymin, ymax). Since the vector data (polygons and points) have inverted y values relative to the image, the spatial selection was first flipped across the image y midline and then used to select the data. The resulting data was then flipped back across the image’s midline and then ingested as `giottoPolygon` and `giottoPoints` objects respectively. Next, Giotto objects were then constructed from the `giottoPoints`, `giottoPolygon`, and `giottoLargeImage` objects using `createGiottoObject()`. For the Breast Cancer dataset, only 1 of the polygon layers was loaded in since they are all identical. For the Mouse Brain dataset, all 7 of the polygon z-layers were loaded. Next, `addSpatialCentroidLocations()` was used to calculate a set of spatial location coordinates for each of the polygons. Counts matrices were then created for each set of polygons in each dataset by first running `calculateOverlapRaster()` to determine the points data overlapped by the polygons, and then `overlapToMatrix()` to convert the overlaps information into matrices. At this step, `aggregateStacks()` was run on the Mouse Brain dataset to combine the expression and spatial information content of its 7 z-layers into a single spatial unit called “aggregate”. This step also creates a new set of matching “aggregate” polygon information, with vertices defined as the combined outer boundary across layers. From here, both datasets were processed using the standard steps, including filtering, normalization, dimension reduction, and clustering.

#### Multiscale analysis and visualization

To demonstrate data representation and analysis at multiple spatial scales, we started with a subset of the Vizgen FFPE Breast Cancer dataset which already had a set of machine-based cell boundary segmentations. We added a set of nuclear segmentations by hand using *QuPath*. Additional spatial scales or spatial units for cell neighborhoods and domains were calculated based on cell-level expression information. Cell neighborhoods were defined by finding local niches composed of cells with similar Leiden cluster results using `calculateSpatCellMetadataProportions()` and kmeans (k = 6). Spatial domains were detected using Hidden Markov Random Field (HMRF)^1^ on genes selected from spatial co-expression modules. `spatialSplitCluster()` was then run to further split the resulting annotations when regions were not spatially touching. Sankey plots were used to show spatial intersections across the previously described spatial units.

#### Transcript location gene set enrichment analysis

Comparative analysis of transcript abundance differences within the cell versus extracellular and nuclear versus cytoplasmic segmented compartments was performed to demonstrate spatial querying. A cell was defined as all transcript detections that overlap within the Vizgen-provided cell segmentation annotations and extracellular as those that are outside. Nuclear was the detections that were within the manual nuclear annotations, and cytoplasmic were those that were within the cell, but outside the nuclear. GSEA analysis was performed using *clusterProfiler*, *biomaRt*, and *org.Hs.eg.db* packages. The results were then plotted using *enrichplot* package’s `dotplot()` function.

#### Z-stack variance analysis

To illustrate variance in transcript abundance between different z-stack layers a subset of the Vizgen Mouse Brain Receptor Map dataset was used. The polygons from each of the 7 provided z-layers were rasterized at 1000 x 1000 with the `rast()` function from *terra*. To assess gene expression variance within and between z-stack layers, transcript locations for each gene were first rasterized and summed within pixels at a predetermined pixel size (20 x 20). Hence, each gene is represented by 7 aligned images and these images were then used to calculate the coefficient of variation (COV) across the z-stack layers for each gene. To identify genes that display higher than expected variation the COV was plotted against the summed counts.

### 10X Genomics data processing and analysis

To demonstrate the different steps of co-registering spatial multi-modal datasets, the data associated with the 10X Genomics Breast Cancer dataset was used. This Xenium instrument data includes subcellular transcript locations, polygon information, immunofluorescence (IF), and post-Xenium H&E images from the same tissue section. From an adjacent tissue section, the Visium CytAssist instrument was used to generate gene expression data with an H&E image.

#### Co-registering multimodal dataset

Image registration was performed using *STalign*^2^. Depending on data input and registration purposes we used both affine and non-rigid transformation modes from *STalign* as explained in more detail below. First, to register Xenium transcripts to the post-Xenium H&E image, we calculated an affine transformation matrix (`Affine_tx`). The corresponding cell centroids were used as input to the `STalign.rasterize()` function, supplying the argument `dx = 15`. This function returned rasterized cell centroid data for landmark detection. Landmark points were manually selected with the point_annotator.py script provided by *STalign*, and this information was provided to the function `STalign.L_T_from_points()` to calculate the affine matrix, `Affine_tx`. The coordinates of cell boundaries were also applied with the same `Affine_tx` matrix along with the registration of transcripts. Next, to register the IF image to the post-Xenium H&E image, the IF image data was converted to point data, represented as pixel intensity, x, and y coordinates. The affine transformation matrix for the IF data (`Affine_IF`) was then calculated following the same pipeline for the creation of `Affine_tx`. Then, large deformation diffeomorphic metric mapping (LDDMM) was performed to register Xenium data to the Visium H&E image. The affine matrix for Xenium to Visium H&E registration (`Affine_visium`) was first calculated in the same manner described above. Default parameters were chosen to optimize the velocity field with `STalign.LDDMM()`. The combined deformation field (`Φ_visium`) was applied to Xenium transcript locations to complete the registration. In parallel, the polygons within the Visium coordinate system that represent Visium spots were also mapped to the Xenium H&E image by serially calculating the inverse operation of `Affine_visium`. This was done by swapping the input order of landmarks for the function `STalign.L_T_from_points()`, and then transforming with `Affine_tx` so that the spatial locations of Visium counts were registered to the same coordinate system of the post-Xenium H&E image. Finally, the IF, Xenium transcript locations, corresponding segmentation polygons, and Visium counts were aligned to the post-Xenium H&E image coordinate system. This resulted in a fully aligned and multi-modal dataset with overlapping regions for all three modalities.

#### Cell segmentation

The 10X Genomics Xenium Pre-Release Breast Cancer dataset was segmented with different methods to simulate different scenarios or allow method comparison within the Giotto Suite framework. The original segmentation data is stored in .csv data format and can be directly read into a `giottoPolygon` object using the `createGiottoPolygons()` function. Other segmentation methods were optimized by following their instructions and exploring their parameter space. A final segmentation result for each method was selected based on empirical observation of the resulting polygons and details are described below.

*Baysor*^3^ (version 0.5.2 with Julia version 1.7.3) was used in combination with a reference segmentation, i.e. the original segmentation results provided by 10X genomics for initialization. The minimum transcripts per cell parameter was set to 30, and the confidence for the prior segmentation was set to 0.5. The resulting *Baysor* polygons were saved as a `geoJSON` file, converted to a data frame with polygon vertices, and used as input for the `createGiottoPolygonsFromDfr()` function.

*CellPose*^4^ (v2.2.2) was used to segment the Xenium IF data. The IF image was segmented in Python using the `cyto2` model from *CellPose* out-of-the-box. The HER2 channel was defined as a cell boundary stain for the model, while the DAPI channel was used as the nuclei stain. No further parameters were modified. The mask output was written to a tif file using the cv2 library and provided as input to the `createGiottoPolygonsFromMask()` function.

The IF image was loaded into *QuPath*^5^ (v0.4.3) and segmented using *Universal StarDist for QuPath*^6^. StarDist^7^ was installed as a *QuPath* extension (link). Pretrained models were downloaded from a repository (link), and the model ‘dsb2018_heavy_augment.pb ’ was used in the final analysis. The file ‘GPU_Multimodal StarDist Segmentation.groovy ’ was downloaded from a separate repository, and was used as the segmentation script on QuPath. The following parameters were used in the previous file: model_trained_on_single_channel = 1, param_channel = 3, param_median = 0, param_divide = 1, param_add = 0, param_threshold = 0.5, param_pixelsize = 0, param_tilesize = 768, param_expansion = 10, min_nuc_area = 10, max_nuc_area = 1000, nuc_area_measurement = “Area px^2”, min_nuc_intensity = 0.5, nuc_intensity_measurement = “DAPI: Nucleus: Mean”, normalize_low_pct = 1, normalize_high_pct = 99. The script was adjusted to remove detections of nuclei with a total area larger than the maximum nuclei size parameter. The resulting segmentation data was then exported from *QuPath* as a geoJSON file and provided as input to `createGiottoPolygonsFromGeoJSON()`.

#### Spatial correlation analysis of molecular modalities from adjacent tissue slices

*Transcript and Protein Colocalization Analysis.* A subcellular Giotto Object was created using the transcript locations which had been aligned to the corresponding Xenium H&E image, as previously described. The immunofluorescence data from the Xenium experiment was aligned, as previously described, and split into single-channel images. The data for the HER2 and CD20 channels were separately rasterized using *terra*. Each of these rasters was then converted to a `giottoLargeImage` and added to a Giotto Object. The transcript locations of ERBB2 and MS4A1 were extracted from the Giotto object and each was rasterized in the same manner. The corresponding `spatRaster` objects were combined (i.e. HER2 and ERBB2), and zero values were imputed for any resulting NAs. The combined raster objects were each provided to the *terra* function `layerCor()` to calculate a simple Pearson’s correlation between the transcript counts and intensity information.

*Xenium aggregated transcripts and Visium.* The Xenium transcript data was registered to Visium H&E Image using the deformation field `Φ_visium` as described above. After loading the registered Xenium data in Giotto Suite pseudo-Visium polygons were created at the same location of the spots from the Visium data using the function `polyStamp()`. An aggregated transcript matrix within each pseudo-spot was created with the functions `calculateOverlapRaster()` and `overlapToMatrix()`, which identifies overlapping transcript and polygon coordinates and converts the overlapping results into a matrix, respectively. The Pearson correlation statistic was then calculated using the log10-normalized values for all common genes between Visium and Xenium. To assess the association between correlation scores and gene expression levels, we computed the Loess trend for the log10 expression value of both Xenium and Visium assays against the rank of the correlation of intersecting genes.

#### Joint clustering and comparing of multiple segmentation results

*Creation and processing of a Multi-Segmentation Giotto object.* Two subcellular Giotto objects were created using the 10X Genomics Xenium Pre-Release Breast Cancer dataset. The first Giotto object was created using the original cell segmentation data, while the second was created using the aligned `Baysor` cell segmentation data. For this analysis segmentation data was provided in a .csv format which is the output format of the *STAlign* co-registration pipeline that was used. The gene-by-cell expression matrix for each Giotto Object was calculated using the functions `calculateOverlapRaster()` and `overlapToMatrix()`. Cell centroids were then computed from the segmentation polygons using the `addSpatialCentroidLocations()` function and used as spatial locations in downstream analyses. Next, the two Giotto objects were combined using the function `joinGiottoObjects()`, which is typically used for analyzing multiple samples by appending sample-specific names (e.g. baysor) to cell IDs from both objects. The combined Giotto object was then processed following standard processing steps, including filtering, normalization, and dimension reduction with PCA.

*Clustering and annotation.* A downsampled Giotto object was created using 25% of all cells (n = 96,329) and used in downstream Giotto projection functions to speed up exploratory data analysis and cell clustering. First, this subset was used to create a low-dimensional UMAP representation (`runUMAPprojection())` and nearest neighbor network (`createNearestNetwork())` using the top 25 PCs. Leiden clustering was then performed with the `doLeidenClusterIgraph()` function, with 100 iterations and a resolution parameter of 0.55. The obtained Leiden cluster labels were then mapped back to the larger, joint Giotto object through the use of `doClusterProjection()`, which uses a knn classifier to assign a label to each unseen cell by majority vote on the same PCA space (top 25 PC), with ties broken at random. Finally, individual Leiden clusters from the whole Giotto object were annotated based on differential gene expression information (i.e. known markers) and visual overlap with the original annotations provided by 10X Genomics.

*Pairwise cluster results comparison.* The average polygon size of each segmentation method was calculated by averaging the results of the `expanse()` function from *terra*, with the segmentation-specific `giottoPolygon` provided as input. Segmentation-specific cell metadata was extracted from the joint Giotto Object by logical indexing of the `list_ID` column. The cell types comprising each segmentation method were identified and put into tabular format to determine occurrence frequency. Percentages of each cell type were calculated by dividing the number of occurrences per cell type by the number of polygons in the segmentation method.

### Imaging Mass Cytometry data processing and analysis

The human lymph node FFPE IMC dataset contains 12 images. First, the 193Ir intensity image, depicting nucleic acids, was used for segmentation in *QuPath*, using the Positive Cell Detection functionality (Non-default parameters: Background Radius:5, Minimum Area:5, Maximum Area:18, Threshold:5, Smooth Boundaries:No). The polygonal data was exported from *QuPath* as a .geojson file. Intensity images corresponding to the following genes were used to create a subcellular Giotto object using `createGiottoObjectSubcellular()`: MX1, Ki67, CD20, CD69, DNA1, Actin, FoxP3, MMP9, CXCL13, CD45, Vimentin. The subcellular locations in the Giotto object were subsetted, and centroids were calculated for all polygons within the default spatial unit “cell”.

#### Polygon rescaling analysis

The function `rescalePolygons()` was used to create a new set of polygons stored in a separate spatial unit, “smallcell” (Arguments used: `poly_info = “cell”, name = “smallcell”, fx = 0.75, fy = 0.75, calculate_centroids = TRUE`). For both spatial units, overlaps between polygon and feature information (i.e. intensities for each antibody representing the genes previously listed) were calculated using `calculateOverlapPolygonImages()`. The features were aggregated by summation per polygon to create an expression matrix using `overlapImagesToMatrix()`. Using the function `filterGiotto()`, both spatial units in the Giotto object were filtered using the same parameters (`expression_threshold = 0, feat_det_in_min_cells = 10, min_det_feats_per_cell = 2`). The expression threshold was left at 0 in order to preserve the maximum amount of expression information. Expression matrices for each spatial unit were normalized with `normalizeGiotto()` via the pearson_resid method, and PCA dimension reductions were performed using `runPCA()` on the sets of normalized values (`scale_unit = FALSE, center = FALSE, ncp = 20`). UMAPs were calculated with `runUMAP()` for each spatial unit (`dimensions_to_use = 1:5`), and a shared nearest network was created with the `createNearestNetwork()` function (`dimensions_to_use = 1:5`). Kmeans clustering was performed on the normalized values of each spatial unit with `doKmeans() and k = 7` for each spatial unit. Between the spatial units, clusters were determined to be corresponding based on the percentage overlap in cell_IDs in each cluster. Clusters with maximal overlapping cell IDs were determined to be matching clusters between spatial units “cell” and “smallcell”. This was calculated with the convenience function `showPolygonSizeInfluence()`. The results were visualized with `spatInSituPlotPoints()`.

### Multi-omics Visium CytAssist data processing and analysis

#### Multi-modal integration

To perform multi-modal data integration, individual modality PCAs and k-Nearest Neighbors were used for calculating the weighted matrix and cell-specific modality weights using the method of Hao, *et al* integrated into the function `runWNN()`. Both results were stored within the multi-omics slot of the Giotto object. The resulting weighted matrix was used for calculating an integrated k-Nearest Neighbor graph by using the function `runIntegratedUMAP()`. Both weighted matrix and integrated kNN graph were used for calculating the integrated UMAP that was stored within the dimension reduction slot of the Giotto Suite object, and using both RNA and protein feature type names. Finally, the integrated kNN graph was used to calculate integrated Leiden clusters using the standard Giotto function `doLeidenCluster()`. The resulting cluster IDs were stored in the cell metadata slot.

#### Analysis of Human Tonsil dataset

For RNA modality, a minimum of 1000 features per cell, 50 cells with a feature, and an expression threshold of 1 was used for filtering, resulting in the removal of 3 out of 4194 cells and 230 out of 18977 genes. For the Protein modality, a minimum of 1 feature per cell, 50 cells with a feature, and an expression threshold of 1 were used, none cells nor proteins were removed. Both RNA and protein expression matrices were normalized and scaled using a `logbase = 2`, `log_offset = 1`, and a `scalefactor = 6000`. The calculation of highly variable features was performed for RNA modality using a Z-score `threshold = 1.5`. The HVFs were used for the calculation of the RNA modality PCA, while all 35 proteins were used for protein PCA; 100 Principal Components were calculated for both individual modalities. The first 10 PCs were used for the calculation of individual UMAP, tSNE, and shared nearest neighbors. The resulting sNN graphs were used for calculating Leiden Clusters with a resolution of 1. The individual modality PCAs were used for integrating RNA and protein modalities, and then the integrated UMAP and clusters were calculated by running Giotto functions `runWNN()`, `runIntegratedUMAP()`, and `doLeidenClusters()` respectively. The annotated single-cell dataset from the Atlas of Cells in the Human Tonsil at the annotation level 1 and the integrated Leiden clusters were used for the deconvolution of the ST dataset using the *SpatialDWLS* method.

### Stereo-seq data processing and analysis

Bin1 matrix files were downloaded from the CNGB website (see Data and Code Availability for details), unzipped using gunzip, and converted into .bgef files using `generate_bgef()` from the *gefpy* python module (v0.5.4.8). A reader function using the *rhdf5* package was written to combine the .bgef `/geneExp/bin1/expression` and `/geneExp/bin1/gene` datasets into gene detections per bin. These values were ingested chunkwise into a *duckdb* backend as a `dbPointsProxy` using `dbvect()`. `tesselate()` was then used to generate two sets of tiled hexagon bin polygons, with diameter of 400 and 100 units (200 and 50 microns respectively), which were also read into the backend as `dbPolygonProxy` using `dbvect()`. Finally, `calculateOverlap()` was run on the `dbPolygonProxy` and `dbPointsProxy` objects and the overlap results were written to disk using `overlapToMatrix()` as HDF5Matrix count matrices. The matrices and the tesselated polygons were added to a Giotto Object. The expression information was filtered and normalized with `filterGiotto()` and `normalizeGiotto()`, then `calculateHVF()` was used to find highly variable genes using a randomly sampled 10% subset of the dataset. Downstream, further projection strategies were used to speed up analysis. `runPCAprojection()`, `runUMAPprojection()`, and `doClusterProjection()` were performed with 25% subsets of the dataset, after which the results were projected onto the rest of the dataset. See the script for further details and parameters.

#### Resolution increase, pseudo-aggregation and deconvolution

A representative region of interest was selected in the mouse brain with `subsetGiottoLocs()` and coordinates x_min = 1000, x_max = 3000, y_min = 9000, and y_max = 12000. Next, a similar workflow as for the whole embryo dataset was followed, except with hexagon bin polygons with diameter of 50 units (25 µm). In addition, the Leiden cluster results were used to perform niche clustering with `calculateSpatCellMetadataProportions(`) and simple `kmeans()` (k = 12). To identify spatial co-expression modules a spatial k-nearest neighbor network was created with `createSpatialNetwork()` and parameters k = 10 and maximum_distance_knn = 200. This network was used together with binSpect() to identify the top 500 spatial genes starting from all highly variable genes. These most spatially variable genes were used as input for detectSpatialCorFeats() to compute spatial co-expression modules and followed by hierarchical clustering with clusterSpatialCorFeats() and k = 10. Individual spatial co-expression modules were converted to metafeatures (i.e. metagenes) with `createMetafeats()` and visualized with `spatCellPlot()`. Next, `makePseudoVisium()` was used to generate a scale-accurate Visium spot array within the region of interest. For deconvolution, spatialDWLS was run using `runDWLSDeconv()` on the pseudo-Visium spots using a previously published developmental mouse brain single-cell atlas as a reference dataset for cell typing with the blood cell type removed (see Data and Code Availability for sources). Top single cell markers in this dataset were identified using the `findMarkers_one_vs_all()` function in Giotto Suite using the scran method.

### Scalability implementations

#### Integration of Giotto in the Cloud

We developed a Docker image compatible with terra.bio available at giottopackage/terra_jupyter_suite_modular:latest. The image contains the latest version of Giotto 3.4.0 which allows running interactive Jupyter notebooks within a customized cloud environment. On the other hand, we developed a startup script for running the RStudio app with an automatic Giotto installation, available at https://github.com/drieslab/Giotto_Suite_manuscript.

#### DelayedArray and future.apply Implementation

We used the HDF5Array package to create a DelayedArray backend. For integrating the HDF5 backend, the function `createGiottoObject()` was adapted to write expression matrices within an on-disk .h5 file instead of the Giotto object, while a string with the internal path in the h5 file leading to the matrix was stored in the expression slot. Giotto getter and setter functions were adapted to automatically identify the HDF5 backend and manage expression information to/from the on-disk file using the *chihaya* package. Additionally, the *ScaledMatrix* package was used for storing and reading scaled matrices. Analysis functions were adapted using the *DelayedMatrixStats* package to handle *DelayedArray* calculations. To allow parallel operations the *future.apply* package has been implemented and users can follow the `plan`() guidelines to use the processing (e.g. sequential or multisession) backend of choice.

#### Subsampling and projection strategies

To facilitate large-scale principal component analysis, `runPCAprojection()` and `runPCAprojectionBatch()` were implemented. First, the expression matrix is subsetted by taking a user-defined percentage of all spatial units (e.g. cells) in a random sampling manner. Next, the downscaled expression matrix is used for PCA using the standard implementations in Giotto Suite and results are converted to an S3 *prcomp* class. This is then followed by the projection of the remaining expression matrix with `predict.prcomp()` to the same PCA space. A similar approach is followed by the batch approach, except that multiple batches will be performed and aggregated for a final PCA result. To compute UMAP coordinates from a large-scale spatial dataset `runUMAPprojection()` was implemented. First, the expression matrix is subsetted by taking a user-defined percentage of all spatial units (e.g. cells) in a random sampling manner. Next, the downscaled expression matrix is used with `runUMAP()` and the UMAP model is stored. This UMAP model is subsequently used to transform the remaining expression matrix spatial units (e.g. cells) into the same UMAP coordinate space. Finally, `doClusterProjection()` is implemented to transfer annotation labels from spatial units (e.g. cells) to other unseen spatial units that share an identical dimension reduction space. For this approach, users can create a smaller Giotto object with one of the convenience functions (e.g. `subsetGiotto()`), cluster the data with their preferred method (e.g. kmeans, hierarchical, Leiden, etc), and subsequently provide both the original Giotto object (target) and the smaller cluster Giotto object (source) to transfer the obtained labels using a fast k-nearest neighbor approach as implemented by the *FNN* package.

#### Database and spatial chunking approach

`dbPointsProxy` and `dbPolygonProxy` are S4 structures that contain *dbplyr*/*dplyr* `tbls` connected to a database via *DBI*. They can be created using `dbvect()` from specifically formatted `data.frames`, *terra* `SpatVectors,` and filepath inputs. In these analyses, the database used was *DuckDB*. On backend creation, connection details are stored in a package-level environment, from which objects can independently retrieve connections, allowing them to function in a standalone manner and be encapsulated within larger objects similarly to normal in-memory objects. Connection handling for these objects is then abstracted away through *pool*. These representations respond to spatial manipulation generics and can be pulled into memory as their corresponding *terra* objects using `as.spatvector()`, making them convenient proxies for the data that they contain. Spatially chunked processing is implemented through `chunkSpatApply(),` which plans and pulls out chunks of data for up to two inputs and then applies a supplied function, writing the results back to the database. Individual geometries are selected during the chunking process using a *min value* ≤ *x* < *max value* filter, ensuring entities are not double-selected. For `dbPolygonProxy` specifically, geometries are selected based on the x and y means of the vertices of each polygon. Utilities for table generation with constraints, and chunkwise data ingestion into the database are also provided and allow flexible use of different reader, writer, and callback functions.

### Embedded Shiny Interactivity

#### Interactive polygon selection

We developed a *Shiny* gadget that launches a local application to interactively draw multiple regions of interest over a Giotto spatial plot, by running the function `plotInteractivePolygons().` The spatial plot may or not contain a tissue image in the background. The application provides the flexibility to assign custom names for each region of interest, as well as multiple or individual colors for the polygons. The tool also provides slide bars across the x and y axes to zoom -in and -out over the image. The reactivity feature of this interactive plot allows users to draw new polygons on the images, as well as simultaneously retrieve the corresponding x and y coordinates to a user-defined variable within the R console. The resulting table with coordinates can subsequently be used or integrated within the Giotto Suite object by running the functions `addGiottoPolygons()` and `addPolygonCells().` Polygon information can be used for downstream analysis such as the comparison of cell type abundance or gene expression patterns within the drawn areas by running the functions `compareCellAbundance()` and `comparePolygonExpression()`, respectively.

#### Interactive 3D spatial plotting

To create an interactive tri-dimensional visualization of 3D spatial datasets the *plotly* package was used within an interactive *Shiny* application. The implementation runs locally by calling the function `plotInteractive3D()`. The application is reactive to slide bars that modify the lower and upper limits of the x, y, z axis creating custom slices across the dataset. Additionally, the plot is reactive to an optional selection of cluster IDs listed in the cell metadata table, facilitating the visualization and subsetting of cell types of interest. When closing the application, a table containing cell IDs, spatial coordinates, and cluster or cell type IDs will be retrieved.

### Interoperability

#### Converters between Giotto Suite and other spatial omics packages

Currently, Giotto objects created within Giotto Suite are interoperable with other spatial omics packages, including *Bioconductor/SpatialExperiment*, *Seurat*, and *AnnData/Squidpy*. This promotes a bi-directional compatibility of Giotto objects with other ecosystems and simultaneously extends its applications.

For the Bioconductor group of packages, the *SpatialExperiment* data container is used for storing data from spatial-omics experiments. It is designed to handle data from spot-based and molecule-based platforms that include spatial coordinates, images, and image metadata, apart from the data already common to Experiment classes. Giotto Suite provides two functions `giottoToSpatialExperiment()` and `spatialExperimentToGiotto()` developed by mapping the slots of the Giotto object to the corresponding slots. Briefly, Giotto’s feat_metadata maps to SpatialExperiment’s rowData, expression corresponds to assays, cell_metadata to col_Data, dim_reductions to reducedDims, spatial_locs to spatialCoords and Images are reflected as imgData. The images in Giotto are technically stored as raster objects and *SpatialExperiment* also supports the same. Giotto handles expression matrices within separate spatial units and feature types. The `SpatialExperiment` object can only store one spatial unit at a time therefore, a list of `SpatialExperiment` objects is returned from the `giottoToSpatialExperiment()` function, where each element of the list corresponds to a distinct `SpatialExperiment` object for a specific spatial unit.

Giotto Suite also provides interoperability between *Seurat* and Giotto. Since *Seurat* has multiple versions in use with differences in object structure, we currently provide interoperability between Giotto and both the older and the newer versions of Seurat objects. Therefore, there are four functions tailored for these different Seurat versions: `giottoToSeuratV4()` and `seuratToGiottoV4()` for the older versions, and `giottoToSeuratV5()` and `seuratToGiottoV5()` for Seurat v5, which now includes subcellular and image information. The v4 functions map Giotto’s cell_metadata to Seurat’s meta.data, dimension_reduction to reductions, feat_metadata from Giotto is mapped to meta.data for each assay in *Seurat* and expression to assays. With v5, additional slots like spatial_loc and images from Giotto are mapped to the most relevant slots in *Seurat*. During the conversion from Giotto to *Seurat*, Giotto’s spatial information is stored in Seurat’s dimension reduction slot as it does not provide a separate slot for overall tissue level coordinates. Images and subcellular information in Giotto are both passed to the images slot of the Seurat objects.

Finally, to support conversions to the `AnnData` class in Python, the functions `anndataToGiotto()` and `giottoToAnnData()` were created by mapping the slots of the Giotto object to the corresponding locations in a squidpy-flavored `AnnData` object. In summary, Giotto’s expression slot maps to adata.X, spatial_locs to adata.obsm, cell_metadata to adata.obs, feat_metadata to adata.var, dimension_reduction to adata.obsm, nn_network and spat_network to adata.obsp. Images are currently not mapped between both classes. Of note, the Giotto object stores expression matrices within separate spatial units and feature types, while `AnnData` objects do not support this hierarchical data storage method. Thus multiple `AnnData` objects will be created from a Giotto object when multiple spatial units and feature type pairs exist.

#### Bento integration and analysis

To integrate *Bento* analysis with Giotto, *Bento* (version 2.0.1) scripts were adapted and updated to ensure compatibility with Python 3.10, the current default version utilized in Giotto Suite. These modified *Bento* scripts are also accessible on GitHub at https://github.com/wwang-chcn/bento-tools. For this example, the Xenium Breast Cancer dataset was used and subsetted with `subsetGiottoLocs()` with parameters `x_min = 0, x_max = 2000, y_min = 0, and y_max = 2000`. First, transcripts, cell, and nuclei coordinates information was extracted from the Giotto object. Next, cell and nucleus polygons were re-created in Python utilizing the *shapely* package (v1.8.5.post1) and subsequently stored in a *geopandas* (v0.10.2)) dataframe. A modified `AnnData` object was created using the `bt.io.prepare()` function from *Bento*. Furthermore, various APIs were established to invoke *Bento’s* tools and plotting functions for shape features, point features, RNAflus, RNAforest, and colocalization analyses. amounting to a total of 12 APIs.

#### R Spatial open science integration and analysis

To facilitate integration between Giotto Suite and (geo)spatial open science ecosystems we have implemented several convenience and interoperability functionality. First, functions and methods from the *terra* package are also available for the derived `giottoPoints` and `giottoPolygon` objects. Next, to scale up and generalize accessibility for other (geo)spatial classes and dependent packages, we implemented converters between spatial data and *sp*, *sf*, *terra*, and *stars* objects as `as.sp()`, `as.sf()`, `as.terra()`, and `as.stars()`, respectively. These converters can also be directly applied on `giottoPoints` and `giottoPolygon` objects to change the underlying data representation. Finally, to make all spatial auto-correlation statistics and metrics available the functionality from the *spdep* package is also available through the `spdepAutoCorr()` function.

#### Spatial resolution enhancement through interpolation and single-cell segmentation

*Spatial interpolation for spatial variable genes.* The 10X Genomics Mouse Brain Coronal Section dataset was downloaded and used in this analysis (see Data Availability). `createGiottoVisiumObject()` was used to create a Giotto object, considering only in-tissue spots. Raw data was filtered and normalized. `createSpatialnetwork()` was used to generate a spatial network with the ‘kNN’ method, specifying `k=5` and `maximum_distance_knn=400`. This spatial network was used to identify spatially variable features using `binsSpect()` or `spdepAutoCorr()` which makes all metrics and spatial statistics from the *spdep* package available in Giotto Suite. The top 500 spatially variable features were extracted and clustered to identify spatial patterns using the spatial co-expression pipeline. Next, the expression levels of each of these 500 genes were rasterized into 50 row by 50 column *terra* `spatRaster` objects (2,500 pixels). To increase gene expression resolution a raster of 100 rows by 100 columns (10,000 pixels) was first generated with the `rast()` function from *terra* and ensuring that extents of both rasters match the extent of the underlying polygonal representation of Visium spots. Together, the data was used to create a model using `gstat()`, specifying `formula=count∼1` and `locations=∼x+y`. This model, together with the high-resolution raster, was then provided to the `interpolate()` function from *terra* to compute spatial interpolated and higher resolved raster objects for each gene (hires images).

*Building and representing a super-resolved Visium dataset.* The hires images were first converted to a list of *giottoLargeImage* objects using `createGiottoLargeImage()`. Alternatively, these high-resolution images could also be saved as .tif files. The high-resolution H&E image from the dataset was segmented using *StarDist* for *QuPath*. Polygons with a nucleus area less than 0.0001 px^2^ or more than 600 px^2^ were removed. The polygons were exported from *QuPath* in a `geoJSON` format and read into R as `giottoPolygons` using `createGiottoPolygonsFromGeoJSON()`. The provided scalefactor and spatial location data were used to create polygons to represent the original Visium spots. The list of 500 `giottoLargeImages` and both sets of polygonal information were fed to `createGiottoObjectSubcellular()`. Expression matrices were created for each spatial unit (i.e. Visium Spots and StarDist Cells) using the functions `calculateOverlapPolygonImages()` and `overlapImagesToMatrix()`. Expression data for *StarDist* was filtered to remove all cells that did not have any expression data from the 500 genes considered in this analysis. Then, each of the expression matrices was normalized. PCA was performed on both expression matrices, and the first 10 PCs were used with 25 neighbors to calculate a UMAP and shared nearest neighbor network. Finally, the data was clustered using `kmeans()` (`k = 12)`.

